# Enhanced stress tolerance through reduction of G3BP and suppression of stress granules

**DOI:** 10.1101/2020.02.03.925677

**Authors:** Anna K. Lee, Jonathon Klein, Klementina Fon Tacer, Tessa Lord, Melissa J. Oatley, Jon M. Oatley, Shaina N. Porter, Shondra M. Pruett-Miller, Elena B. Tikhonova, Andrey L. Karamyshev, Peiguo Yang, Hong Joo Kim, J. Paul Taylor, Patrick Ryan Potts

## Abstract

Stress granules (SG) are membrane-less ribonucleoprotein condensates that form in response to various stress stimuli via phase separation. SG act as a protective mechanism to cope with acute stress, but persistent SG have cytotoxic effects that are associated with several age-related diseases. Here, we demonstrate that the testis-specific protein, MAGE-B2, increases cellular stress tolerance by suppressing SG formation through translational inhibition of the key SG nucleator G3BP. MAGE-B2 reduces G3BP protein levels below the critical concentration for phase separation and suppresses SG initiation. Importantly, knockout of the MAGE-B2 mouse ortholog confers hypersensitivity of the male germline to heat stress *in vivo*. Thus, MAGE-B2 provides cytoprotection to maintain mammalian spermatogenesis, a highly thermo-sensitive process that must be preserved throughout reproductive life. These results demonstrate a mechanism that allows for tissue-specific resistance against stress through fine-tuning phase separation and could aid in the development of male fertility therapies.

## INTRODUCTION

Cells regularly encounter a variety of stresses, therefore, one of their largest challenges lies in their ability to respond to and defend against changing conditions. In addition to sensing different insults, cells must select an appropriate course of action in order to engage the proper pathways to ultimately repair, reprogram, or undergo cell death, depending on the type and severity of the damage. Exposure to various stressors, such as oxidative or heat stress, initiate highly coordinated cellular stress response pathways that result in the inhibition of translation, disassembly of polysomes, and reorganization of mRNAs and proteins into stress granules (SG) (Anderson et al., 2015; Kedersha and Anderson, 2002).

SG are conserved, highly dynamic ribonucleoprotein (RNP) condensates that, like other RNP granules, are thought to form through liquid-liquid phase separation (LLPS) via the collective behavior of protein-protein, protein-RNA, and RNA-RNA interactions (Protter and Parker, 2016; Van Treeck and Parker, 2018). A growing body of evidence has linked mutations that increase SG formation or decrease SG clearance to various age-related neurodegenerative diseases (Dobra et al., 2018; Kim et al., 2013; Mackenzie et al., 2017; Molliex et al., 2015; Nedelsky and Taylor, 2019; Patel et al., 2015). Yet, it remains unclear what role SG play during the stress response and how disturbances in SG dynamics might promote disease progression. In fact, SG have been proposed to have both pro-survival and pro-death functions depending on the type and duration of stress; however, the exposure of cells to either acute or chronic stress has been suggested as a determinant between the assembly of protective or harmful SG, respectively (Arimoto et al., 2008; Reineke and Neilson, 2019; Zhang et al., 2019). Therefore, cells with long lifespans, such as stem cells and neurons, may be especially prone to repeated episodes of stress, and thus, particularly susceptible to the potentially toxic effects associated with SG formation. Intriguingly, the male germline in mammals is extremely vulnerable to heat stress, such that minor increases in testicular temperature result in reduced spermatogenesis and increased risk of infertility (Reid et al., 1981; Rockett et al., 2001; Yin et al., 1997). However, whether these and other stress-prone cells or tissues have acquired mechanisms to modulate their sensitivity to stress and SG dynamics remains unclear.

Recent efforts have focused largely on deciphering the principles that drive SG assembly as well as identifying the protein and RNA constituents of SG (Jain et al., 2016; Khong et al., 2017; Markmiller et al., 2018; Namkoong et al., 2018; Souquere et al., 2009; Wheeler et al., 2016). G3BP1 and its paralog, G3BP2, (collectively referred to as G3BP) are the best characterized SG nucleating proteins and have been shown to be critical for SG assembly, where overexpression induces SG formation in the absence of stress and deletion ablates SG in response to arsenite (Kedersha et al., 2016; Reineke et al., 2012; Zhang et al., 2019). Although the formation of biomolecular condensates is thought to be highly dependent on factors that drive LLPS, such as protein concentrations of key nucleators, whether G3BP protein levels dictate the set point for SG assembly and the cellular stress threshold have not been determined. Moreover, the molecular mechanisms regulating G3BP concentration and whether different cell types and disease states fine-tune G3BP concentration to alter stress tolerance remain unknown.

Melanoma antigen (MAGE) genes encode a family of proteins sharing a common MAGE homology domain (MHD) (Lee and Potts, 2017). Following the emergence of eutherian mammals, the MAGE family underwent a rapid expansion from a single *MAGE* in lower eukaryotes to more than 50 genes in humans (Lopez-Sanchez et al., 2007). Most MAGEs, including *MAGE-B2*, are located on the X-chromosome and are classified as cancer-testis antigens (CTAs) given that they are primarily expressed in the testis but are aberrantly expressed in cancers (Chomez et al., 2001; Fon Tacer et al., 2019; Weon and Potts, 2015). A growing body of studies have revealed that MAGEs assemble with E3 RING ubiquitin ligases to form MAGE-RING ligases (MRLs) and act as regulators of ubiquitination in diverse cellular and developmental processes (Doyle et al., 2010; Lee and Potts, 2017). However, the functions of most MAGEs, including MAGE-B2, have not been fully elucidated and their study has been primarily restricted to cancer cells (Peche et al., 2015).

Here, we identify MAGE-B2 as a regulator of the cellular stress response. We demonstrate that MAGE-B2 inhibits SG formation by reducing protein levels of the concentration-dependent SG nucleator, G3BP. Intriguingly, in an unanticipated deviation from prototypical MRLs, MAGE-B2 functions as an RNA binding protein (RBP) that directly binds to the G3BP transcript and inhibits its translation. MAGE-B2 suppresses G3BP translation by displacing the DDX5 RNA helicase which promotes G3BP translation. Importantly, MAGE-B2 expression is restricted to the testis where it maintains stemness of spermatogonial stem cells (SSC). Moreover, mice lacking the MAGE-B2 ortholog exhibit increased sensitivity to heat stress *in vivo* as measured by increased SG assembly, significantly damaged testis histology, and mouse infertility. Together, these results establish that MAGE-B2 protects the highly thermo-sensitive germline from heat stress, suggesting that calibration of G3BP levels and SG formation by MAGE-B2 enhances the cellular stress threshold.

## RESULTS

### MAGE-B2 regulates stress granule dynamics

To investigate the molecular function of MAGE-B2, we first identified MAGE-B2 interactors by tandem affinity purification (TAP) coupled to mass spectrometry (TAP-MS) using HEK293 cells stably expressing either the TAP empty vector or TAP-MAGE-B2. Analysis of the proteins that were present in the TAP-MAGE-B2 purification but absent in the control TAP vector alone, revealed an enrichment for RBPs, particularly those localizing to SG (Figure S1A and Table S1). Intriguingly, a previous report characterizing the human RNA-binding proteome identified MAGE-B2 as an RBP (Trendel et al., 2019). Therefore, we examined whether MAGE-B2 localizes to SG in U2OS cells. Although MAGE-B2 did not co-localize with the critical SG factor, G3BP1, within SG (Figure 1A), depletion of MAGE-B2 by three independent siRNAs resulted in significantly increased SG formation in response to the oxidative stressor, sodium arsenite (Figures 1B-1E). This enhanced SG phenotype was not the result of spontaneous SG formation (Figure S1B) and was not restricted to U2OS cells, as depletion of MAGE-B2 in HCT116 cells also led to increased SG numbers (Figures S1C and S1D). Furthermore, these results were not specific to oxidative stress, as knockdown of MAGE-B2 led to increased SG formation upon ER stress (thapsigargin), heat stress (Figures 1D and 1E), and translation inhibition (rocaglamide A) in a dose-dependent manner (Figures 1F-1H). Consistent with transient MAGE-B2 knockdown, MAGE-B2 knockout by CRISPR-Cas9 also resulted in increased SG formation (Figure S1E) that could be rescued by re-expression of MAGE-B2 (Figures 1I and S1F). Furthermore, overexpression of MAGE-B2 reduced SG formation and overall G3BP1 expression (Figures 1J and S1G). We next examined the impact of MAGE-B2 depletion on SG dynamics by live-cell imaging of U2OS cells stably expressing G3BP1-GFP. Knockdown of MAGE-B2 both enhanced SG assembly and delayed SG disassembly (Figures 1K and S1H-S1K).

**Figure 1.**
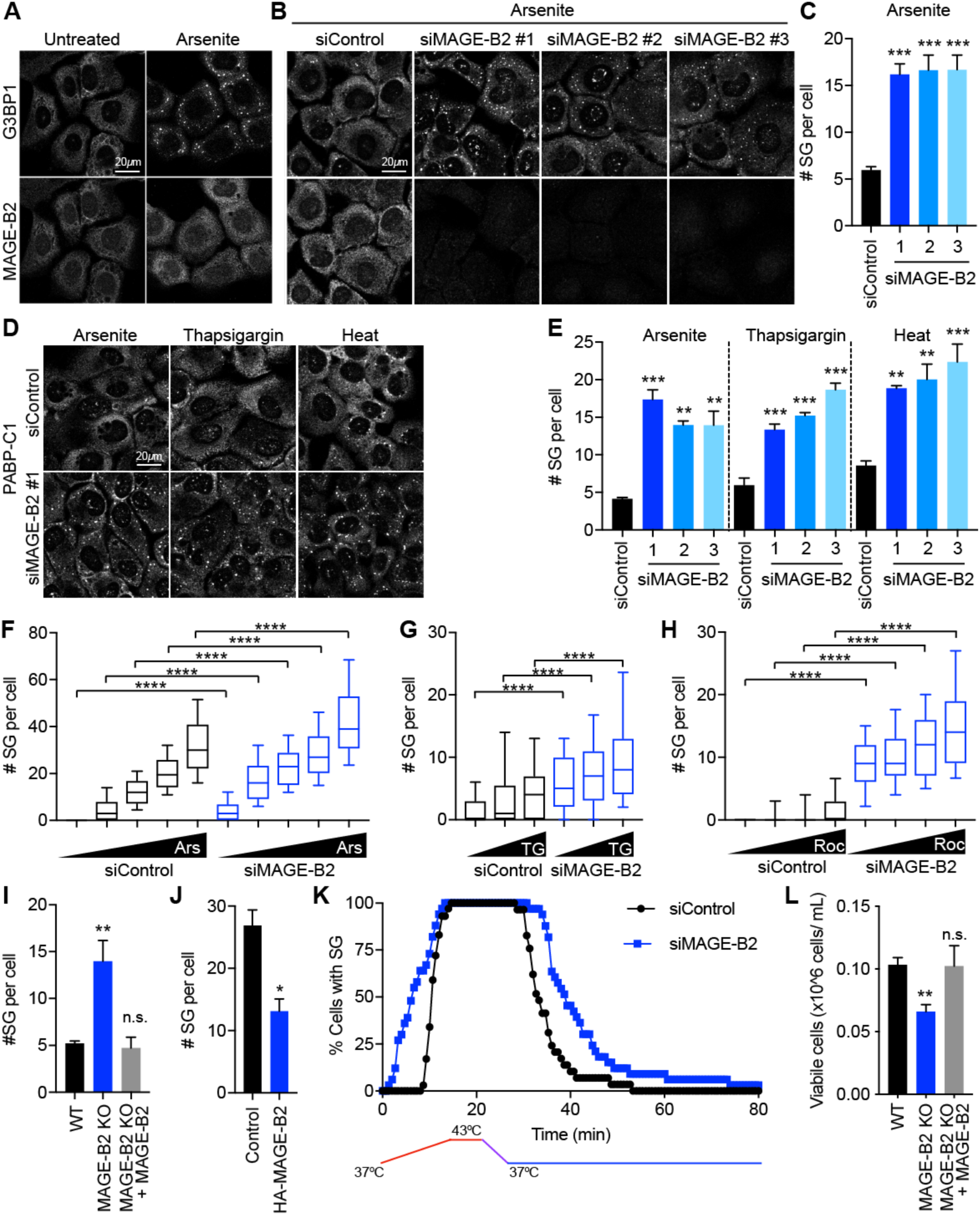
MAGE-B2 regulates SG dynamics. (A) MAGE-B2 does not localize to SG. U2OS cells were treated with or without 500 μM sodium arsenite for 1 hr, and immunostained for G3BP1 and MAGE-B2. Representative images are shown. (B and C) MAGE-B2 knockdown increases SG. U2OS cells were transfected with the indicated siRNAs for 72 hr, treated with 62 μM sodium arsenite for 1 hr, and immunostained for G3BP1 and MAGE-B2. Representative images shown in (B) and quantification (n = 3) of SG numbers per cell shown in (C). *P* values were determined by one-way ANOVA. (D and E) MAGE-B2 knockdown increases SG in response to various stressors. U2OS cells were transfected with the indicated siRNAs for 72 hr, treated with either 62 μM sodium arsenite, 1 μM thapsigargin, or 43 °C heat stress for 1 hr, and immunostained for PABP-C1. Representative images shown in (D) and quantification (n = 3) of SG numbers per cell shown in (E). *P* values were determined by one-way ANOVA. (F-H) MAGE-B2 knockdown increases SG in a dose-dependent manner. U2OS cells were transfected with the indicated siRNAs for 72 hr, treated with either 31 - 500 μM sodium arsenite (F), 0.25 - 1 μM thapsigargin (G), or 0.1 - 1 μM rocaglamide A (H) for 1 hr, and immunostained for G3BP1. Quantification of SG numbers per cell from one representative experiment is shown as a box-and-whisker plot with whiskers drawn down to the 10^th^ percentile and up to the 90^th^ percentile. *P* values were determined by *t* test. WT, MAGE-B2 KO, or MAGE-B2-reconstituted KO U2OS cells were treated with 62 μM sodium arsenite for 1 hr and immunostained for PABP-C1. Quantification (n = 3) of SG numbers per cell is shown. *P* values were determined by *t* test. (I) HA-MAGE-B2 overexpression reduces SG in U2OS cells treated with 500 μM sodium arsenite for 1 hr and immunostained for G3BP1. Quantification (n = 3) of SG numbers per cell is shown. *P* value was determined by *t* test. (J) Live-cell imaging of U2OS cells stably expressing G3BP1-GFP. Cells were heat shocked at 43 °C to induce stress granule assembly and were subsequently recovered at 37 °C to measure stress granule disassembly. Quantification of the percentage of SG-containing cells over time from one representative experiment of n = 3 biological replicates is shown. (K) MAGE-B2 knockout cells are hypersensitive to chronic stress. WT, MAGE-B2 KO, or MAGE-B2-reconstituted KO U2OS cells were exposed to 4 μM sodium arsenite for 3 days before number of viable cells were counted (n = 3). *P* value was determined by *t* test. Data are mean ± SD. Asterisks indicate significant differences from the control (* = p *≤* 0.05, ** = p *≤* 0.01, *** = p *≤* 0.001, **** = p *≤* 0.0001, n.s. = not significant). See also Figure S1.

To determine whether MAGE-B2 regulation of SG dynamics has a functional outcome on cells, we measured the viability of wildtype and MAGE-B2 knockout U2OS cells in response to sodium arsenite. MAGE-B2 knockout cells exhibited hypersensitivity to prolonged low dose sodium arsenite as measured by reduced cell viability that could be rescued by re-expression of MAGE-B2 (Figure 1L). Importantly, wildtype and MAGE-B2 knockout cells grew at similar rates in the absence of stress (Figure S1L). Together, these results suggest that MAGE-B2 inhibits SG formation in response to multiple stressors and that this activity is important for protecting cells against prolonged stress.

### MAGE-B2 modulates SG formation through regulation of G3BP protein levels

Given that MAGE-B2 overexpression led to reduced overall G3BP1 signal by immunostaining (Figure S1G), we hypothesized that MAGE-B2 modulates SG formation by regulating G3BP protein levels. Indeed, depletion of MAGE-B2 by knockdown (Figure 2A) or knockout (Figure 2B) resulted in increased G3BP protein expression. Conversely, overexpression of MAGE-B2 reduced G3BP and re-expression of MAGE-B2 in MAGE-B2 knockout cells rescued G3BP protein levels (Figure 2C). Importantly, these changes in protein expression were specific to G3BP (Figure 2B), such that other SG-associated RBPs were unaffected by MAGE-B2 knockout (Figure S2A). To determine whether MAGE-B2-mediated regulation of G3BP protein levels is responsible for altering SG formation, we rescued G3BP expression in MAGE-B2 depleted cells by concomitantly knocking down G3BP to similar levels as control cells (Figures 2D and S2B). Rescue of G3BP protein back to control levels in MAGE-B2 depleted cells not only rescued the SG phenotype (Figures 2E and 2F), but also cell viability under prolonged stress (Figure 2G). These results suggest that G3BP protein levels are suppressed by MAGE-B2 and this alters the tolerance of cells to stress conditions.

**Figure 2.**
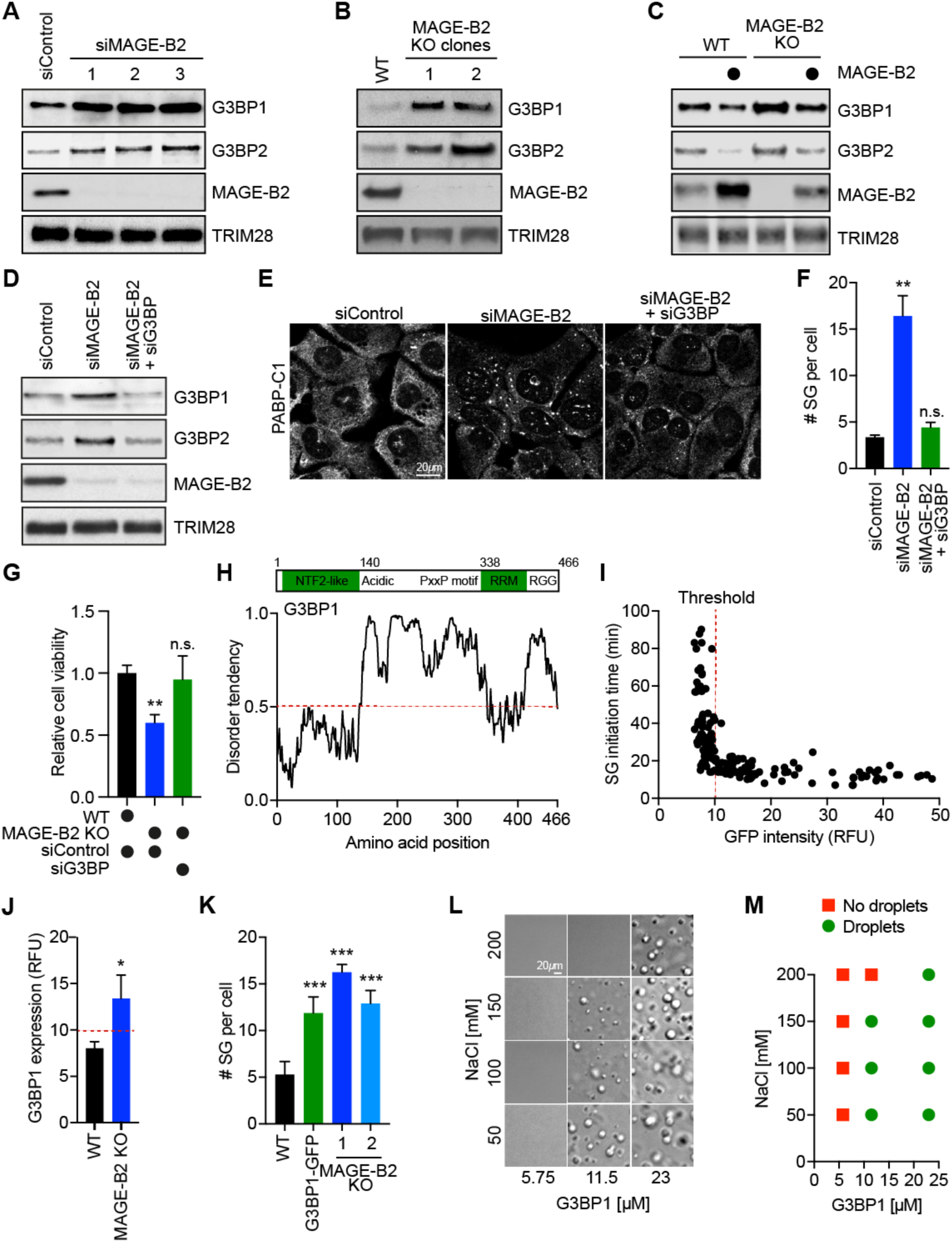
MAGE-B2 modulates SG formation through regulation of G3BP protein levels. (A) MAGE-B2 knockdown increases G3BP protein levels. U2OS cells were transfected with the indicated siRNAs for 72 hr and immunoblotted for the indicated proteins. Immunoblots from one representative experiment of n = 10 biological replicates are shown. (B) MAGE-B2 KO increases G3BP protein levels. WT or MAGE-B2 KO U2OS cell lysates were immunoblotted for the indicated proteins. Immunoblots from one representative experiment of n = 10 biological replicates are shown. (C) Over-expression of MAGE-B2 in WT U2OS cells reduces G3BP protein levels. Stable re-expression of MAGE-B2 in MAGE-B2 KO U2OS cells rescues G3BP protein levels. Cell lysates were immunoblotted for the indicated proteins. Immunoblots from one representative experiment of n = 3 biological replicates are shown. (D-F) Increased G3BP protein levels in MAGE-B2-depleted cells are required for enhanced SG formation. U2OS cells were transfected with the indicated siRNAs for 72 hr and immunoblotted to confirm rescue of G3BP protein levels (D) or treated with 62 μM sodium arsenite for 1 hr and immunostained (E). Quantification (n = 3) of SG number per cell shown in (F). *P* values were determined by *t* test. (G) Depletion of G3BP in MAGE-B2 KO cells restores cell viability during prolonged stress. WT or MAGE-B2 KO U2OS cells were transfected with the indicated siRNAs for 72 hr and exposed to 4 μM sodium arsenite for 3 days before number of viable cells were counted (n = 3). *P* value was determined by *t* test. (H) G3BP1 is an intrinsically disordered protein. Domain map of G3BP1 protein and protein disorder prediction determined are shown. (I) G3BP1 expression correlates with SG initiation time. G3BP KO U2OS cells were transiently transfected with varying levels of GFP-G3BP1. GFP-G3BP1 protein levels were determined at single-cell resolution by live-cell imaging before stress. Cells were then treated with 500 μM sodium arsenite to induce SG assembly and the time at which SG assembly initiated was quantified and correlated to GFP-G3BP1 protein levels (n = 3, total 76 cells analyzed). GFP-G3BP1 expression beyond a threshold (dotted red line) results in enhanced SG formation. (J) MAGE-B2 KO increases G3BP1 protein levels beyond the threshold (dotted red line determined from Fig 2H) for enhanced SG formation. The relative amounts of endogenous G3BP1 in either WT or MAGE-B2 KO U2OS cells was determined in relation to GFP-G3BP1 in Fig 2H (n = 3). *P* value was determined by *t* test. (K) Overexpression of G3BP1 is sufficient to increase SG formation similarly to MAGE-B2 KO. U2OS cells stably overexpressing G3BP1-GFP were treated with 62 μM sodium arsenite for 1 hr, immunostained for PABP-C1, and number of SG per cell was determined (n = 3). *P* values were determined by one-way ANOVA. (L and M) *In vitro* liquid-liquid phase separation of G3BP1. The indicated concentrations of recombinant G3BP1 protein and 100 mg/mL Ficoll 400 were incubated at varying concentrations of NaCl and droplet formation was determined by microscopy. Representative images are shown in (L). Protein/ NaCl concentration pairs scoring positive (green circles) or negative (red squares) for the appearance of droplets from one representative experiment of n = 3 replicates are shown in (M). Data are mean ± SD. Asterisks indicate significant differences from the control (* = p *≤* 0.05, ** = p *≤* 0.01, *** = p *≤* 0.001, n.s. = not significant). See also Figure S2.

### Cellular SG assembly dynamics is dependent on G3BP concentration

Like many other membrane-less organelles, SG are thought to form via LLPS, a biophysical process driven by weak multivalent protein-protein, protein-nucleic acid, and nuclei acid-nucleic acid interactions to create discrete cytoplasmic foci. The crucial molecular features of phase separating proteins include multivalency (tandem arrays of modular domains) and intrinsically disordered regions (IDR), like those found in G3BP (Figure 2H). These proteins phase separate into liquid droplets *in vitro* when their total concentration exceeds a critical threshold, and they return back to a one-phase state once the total concentration drops below the threshold.

Because our results suggest that G3BP protein levels dictate SG dynamics and G3BP has been shown to be a key scaffold for SG assembly, we hypothesized that even modest alterations in G3BP protein levels would contribute to significant changes in SG formation. In order to precisely determine the correlation between G3BP expression and SG initiation time, we transfected G3BP knockout U2OS cells with varying levels of GFP-G3BP1 and determined SG initiation time. Interestingly, the relationship between G3BP1 protein levels and SG kinetics was not linear. Rather, the relationship was switch-like, such that cells expressing G3BP1 beyond the critical threshold exhibited enhanced SG assembly (Figures 2I and S2C), which is consistent with the concept that phase separation is dictated by a critical threshold concentration. Examination of endogenous G3BP1 protein concentrations in wildtype cells revealed that G3BP1 levels are normally maintained just below the threshold (Figures 2J and S2D). However, MAGE-B2 knockout significantly increased G3BP1 beyond the threshold (Figures 2J and S2D). In line with these findings, overexpression of G3BP1 was sufficient to increase SG formation similarly to MAGE-B2 knockout (Figures 2K and S2E). Furthermore, *in vitro* phase separation assays confirmed that small (two-fold) changes in G3BP1 concentration, similar to those observed upon MAGE-B2 knockout, have dramatic effects on G3BP1 LLPS (Figures 2L and 2M). Together, these results suggest that similar to *in vitro* findings, biomolecular condensates in cells are highly dependent on protein concentration; moreover, cells have evolved mechanisms to precisely modulate condensate assembly by directly controlling the key proteins that drive their formation.

### MAGE-B2 downregulates G3BP by translational repression

To determine the mechanism by which MAGE-B2 decreases G3BP protein levels, we tested three potential modes of regulation: transcription, protein stability, and translation. We found that MAGE-B2 did not affect *G3BP1* transcript levels in U2OS, HCT116, or HeLa cells (Figures 3A, S3A and S3B). Our lab and others have previously demonstrated that MAGEs assemble with E3 RING ubiquitin ligases to form MAGE-RING ligases (MRLs) and act as regulators of ubiquitination by modulating ligase activity, substrate specificity, or subcellular localization (Doyle et al., 2010; Hao et al., 2013; Lee and Potts, 2017; Pineda et al., 2015). Therefore, we hypothesized that MAGE-B2 might promote degradation of G3BP. Unexpectedly, G3BP1 protein half-life as measured by ^35^S-methionine/cysteine pulse-chase was unaffected by MAGE-B2 knockout (Figure 3B). In addition, proteasomal inhibition by MG132 in U2OS or HeLa cells did not alter G3BP1 protein levels (Figures S3C and S3D, respectively), consistent with its relatively long protein half-life (∼12 hrs; Figure 3B). Together, these data suggest that MAGE-B2 does not regulate G3BP1 protein levels via the ubiquitin-proteasome system.

**Figure 3.**
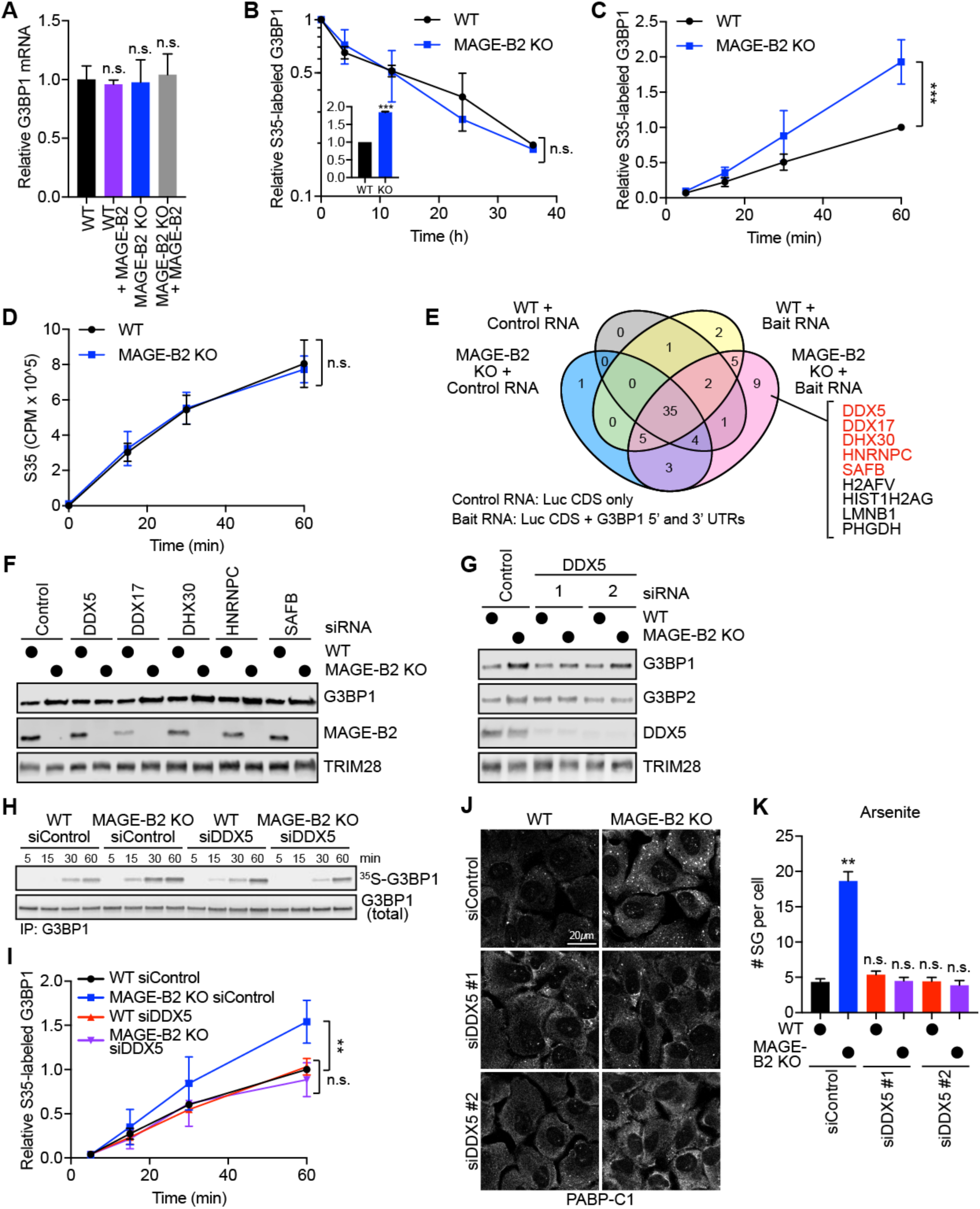
MAGE-B2 represses G3BP1 translation through inhibition of DDX5. (A) MAGE-B2 does not affect *G3BP1* mRNA levels. RT-qPCR analysis (n = 3) of *G3BP1* mRNA levels normalized to *18S* rRNA in the indicated U2OS cells. *P* values were determined by one-way ANOVA. (B) MAGE-B2 does not affect G3BP1 protein degradation. G3BP1 protein half-life was measured (n = 3) in WT or MAGE-B2 KO U2OS cells by ^35^S pulse-chase. Cells were pulse labeled with ^35^S-Met/Cys for 1 hr (t = 0) and then chased in cold Met/Cys for the indicated times. ^35^S-labeled G3BP1 was determined by immunoprecipitation, SDS-PAGE, and autoradiography. *P* value was determined by two-way ANOVA. The inset shows ^35^S-labeled G3BP1 protein levels at t = 0 suggesting differences in G3BP1 translation. *P* value for t = 0 inset data was determined by unpaired *t* test. (C) MAGE-B2 knockout enhances G3BP1 translation. WT or MAGE-B2 KO cells were incubated with ^35^S-Met/Cys for the indicated times before G3BP1 was immunoprecipitated from cell lysates, separated by SDS-PAGE, and the newly synthesized G3BP1 was quantified (n = 3) by autoradiography. *P* value was determined by two-way ANOVA. (D) MAGE-B2 does not affect global translation. Global translation was measured in WT or MAGE-B2 KO U2OS cells by ^35^S-Met/Cys labeling for the indicated times and scintillation counting (n = 3). *P* value was determined by two-way ANOVA. (E) Affinity pulldown of biotinylated RNA. Biotinylated control or bait transcripts (Luciferase CDS with *G3BP1* 5’ and 3’ UTRs) were pulled down from either WT or MAGE-B2 KO U2OS cells and subjected to mass spectrometry analysis. The Venn diagram lists the number of unique proteins for each sample. RNA-binding proteins that bound specifically to the bait transcript in MAGE-B2 KO cells are shown in red. (F) DDX5 knockdown in MAGE-B2 KO cells rescues G3BP1 protein levels. RNA-binding proteins identified in Figure 3E were depleted in WT or MAGE-B2 KO U2OS cells by transfection with the indicated siRNAs for 72 hr to screen for potential involvement in MAGE-B2-mediated regulation of G3BP1. Cell lysates were immunoblotted for the indicated proteins. Immunoblots from one representative experiment of n = 3 biological replicates are shown. (G) Validation of DDX5 knockdown by two independent siRNAs. WT or MAGE-B2 KO U2OS cells were transfected with the indicated siRNAs for 72 hr and immunoblotted for the indicated proteins. Immunoblots from one representative experiment of n = 3 biological replicates are shown. (H and I) DDX5 knockdown in MAGE-B2 KO cells rescues G3BP1 translation. WT or MAGE-B2 KO cells were transfected with the indicated siRNAs for 72 hr before G3BP1 protein synthesis was measured by ^35^S labeling as described in Figure 3B. Representative blot (H) and quantitation (I; n = 3) are shown. *P* values were determined by two-way ANOVA. (J and K) DDX5 knockdown in MAGE-B2 KO cells rescues SG formation. WT or MAGE-B2 KO U2OS cells were transfected with the indicated siRNAs for 72 hr, treated with 62 μM sodium arsenite for 1 hr, and immunostained for PABP-C1. Representative images shown in (J), quantification (n = 3) of SG number per cell shown in (K). *P* values were determined by one-way ANOVA. Data are mean ± SD. Asterisks indicate significant differences from the control (** = p *≤* 0.01, *** = p *≤*0.001, n.s. = not significant). See also Figure S3.

Although the G3BP1 protein half-life was unchanged by MAGE-B2, the relative amounts of ^35^S-labeled G3BP1 immediately after 1 h pulse labeling indicated that G3BP1 protein synthesis was enhanced in MAGE-B2 knockout U2OS cells (Figure 3B, inset). Indeed, ^35^S-methionine/cysteine incorporation assays revealed significantly increased G3BP1 translation in MAGE-B2 knockout cells (Figure 3C). Consistent with our finding that the protein levels of other SG-associated RBPs were unaffected by MAGE-B2 knockout (Figure S2A), MAGE-B2-mediated regulation of G3BP1 translation is specific and not the result of altered global translation as determined by global ^35^S-methionine/cysteine incorporation assays or polysome profiling (Figures 3D and S3E). Together, our results suggest that MAGE-B2 regulates cellular G3BP protein concentrations by downregulating its translation.

### DDX5 mediates MAGE-B2-dependent regulation of G3BP and SG formation

To determine how MAGE-B2 regulates G3BP translation, we hypothesized that MAGE-B2 might affect RBPs bound to *G3BP* UTRs, either by recruiting a translational inhibitor or displacing a translational activator. Therefore, we utilized RNA pulldowns and mass spectrometry to unbiasedly identify proteins differentially bound to *G3BP1* mRNA upon knockout of MAGE-B2. RNA pulldowns from wildtype or MAGE-B2 knockout U2OS cells using biotinylated RNAs consisting of the Luciferase coding sequence (*Luc* CDS) flanked by the *G3BP1* UTRs (bait RNA) or the *Luc* CDS alone (control RNA) were performed and analyzed by mass spectrometry. Interestingly, five RBPs (DDX5, DDX17, DHX30, HNRNPC, and SAFB) bound specifically to the bait RNA in MAGE-B2 knockout cells, but not in wildtype cells or to the control *Luc* CDS RNA (Figure 3E). Notably, a previously described regulator of G3BP1 translation, YB1, was not identified in any RNA pulldowns and its depletion did not affect G3BP1 levels (Figure S3F). However, knockdown of the five candidate RBPs, in wildtype or MAGE-B2 knockout U2OS cells revealed that DDX5 depletion specifically decreased G3BP protein levels in MAGE-B2 knockout cells, but not wildtype cells (Figures 3F and 3G). Additionally, DDX5 protein levels were similar in MAGE-B2 wildtype and knockout cells (Figure 3G). DDX5, also referred to as p68, is a DEAD-box RNA helicase that has been shown to have a number of functions including microRNA processing (Dardenne et al., 2014); however knockdown of AGO2 did not alter G3BP1 protein levels (Figure S3G), suggesting that DDX5-mediated regulation of G3BP1 is independent of its role in microRNA processing. Importantly, knockdown of DDX5 in MAGE-B2 knockout U2OS cells rescued both G3BP1 translation (Figures 3H and 3I) and SG formation (Figures 3J and 3K) without affecting global translation (Figure S3H) or *G3BP1* transcript levels (Figure S3I). These data suggest that DDX5 mediates the enhanced G3BP translation and SG phenotypes observed in MAGE-B2-depleted cells.

### MAGE-B2 and DDX5 have opposing roles in the regulation of G3BP1 translation

We found that MAGE-B2 represses G3BP1 translation and in its absence DDX5 promotes G3BP1 translation, suggesting that MAGE-B2 and DDX5 may have opposing, competing roles. To experimentally determine whether MAGE-B2 and DDX5 bind *G3BP1* mRNA in an opposing fashion, we performed CLIP-qPCR. We found that MAGE-B2 and DDX5 interact with the *G3BP1* transcript (Figure 4A-C). However, DDX5 interaction inversely correlated with MAGE-B2 expression, such that expression of MAGE-B2 decreases DDX5 binding to *G3BP1* transcript (Figure 4B), whereas MAGE-B2 knockout increases DDX5 interaction with *G3BP1* transcript (Figure 4C). Furthermore, using RNA pulldown experiments, we found that MAGE-B2 bound *in vitro* transcribed bait *Luc* CDS with *G3BP1* UTRs, but not control *Luc* CDS alone RNA (Figure 4D). Importantly, DDX5 only bound the *G3BP1* RNA in the absence of MAGE-B2 (Figure 4D). Thus, MAGE-B2 and DDX5 bind to the G3BP1 transcript in an opposing fashion.

**Figure 4.**
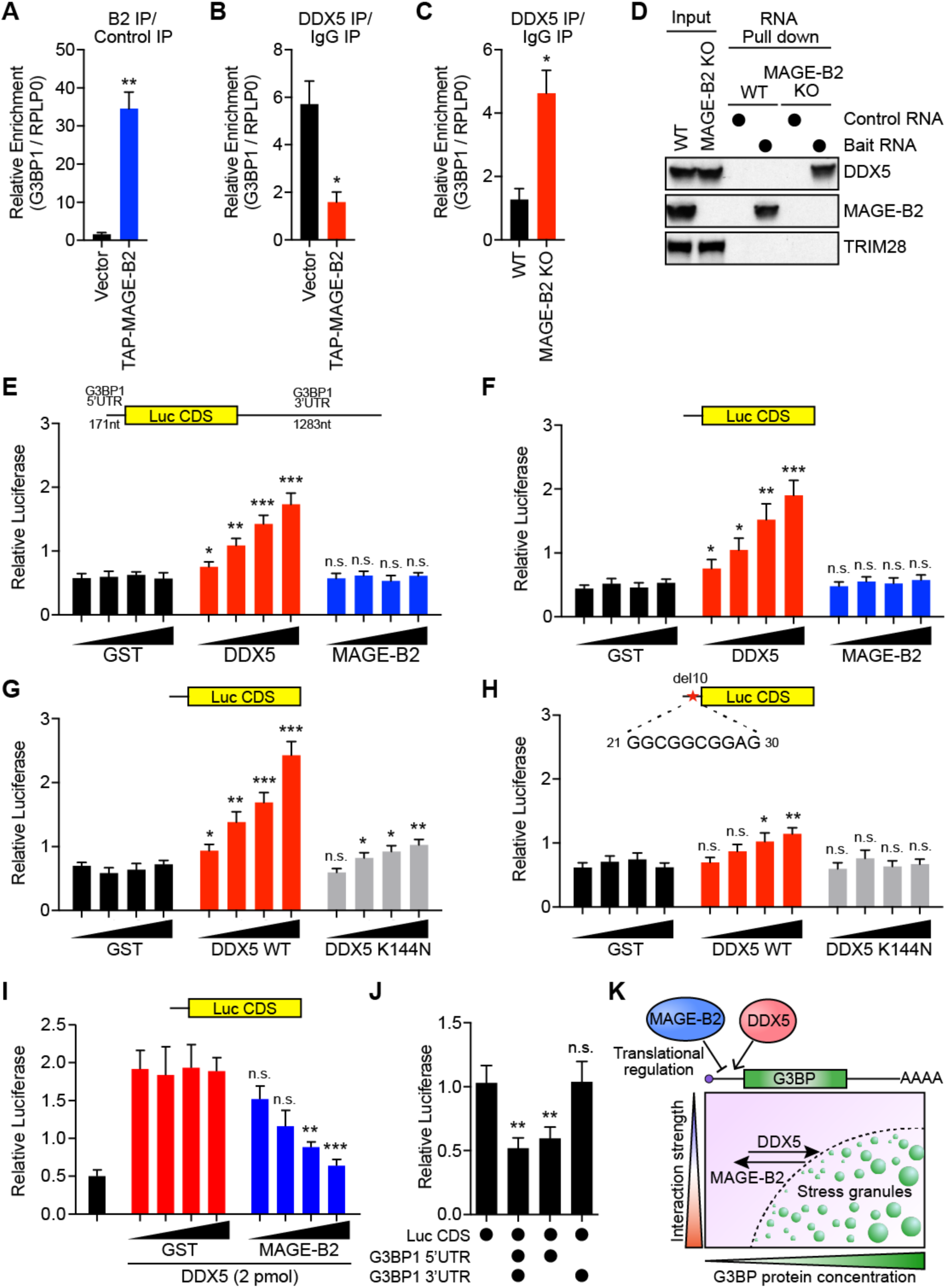
MAGE-B2 and DDX5 have opposing roles in the regulation of G3BP1 translation. (A) MAGE-B2 interacts with *G3BP1* mRNA by CLIP-qPCR. HEK293 cells stably expressing vector control or TAP-MAGE-B2 were subjected to 150 mJ/cm^2^ of UVC (254 nm) before immunoprecipitation of TAP-vector or TAP-MAGE-B2. *G3BP1* mRNA or *RPLP0* mRNA as a normalization control were then detected by RT-qPCR (n = 3). Relative amount of *G3BP1* mRNA in MAGE-B2 pulldown relative to control pulldown after *RPLP0* normalization is shown. *P* value was determined by *t* test. (B) MAGE-B2 suppresses DDX5 interaction with *G3BP1* mRNA. CLIP-qPCR was performed as described above using HEK293 cells stably expressing vector control or TAP-MAGE-B2 to measure enrichment of *G3BP1* mRNA after DDX5 pulldown (n = 3). Relative levels of *G3BP1* mRNA in DDX5 pulldown relative to IgG control after *RPLP0* normalization is shown. *P* value was determined by *t* test. (C) Loss of MAGE-B2 enhances DDX5 interaction with *G3BP1* mRNA. CLIP-qPCR was performed on WT or MAGE-B2 KO U2OS cells to measure enrichment of *G3BP1* mRNA for DDX5 (n = 3) as described above. *P* value was determined by *t* test. (D) MAGE-B2 and DDX5 compete for binding to *G3BP1* mRNA. Biotinylated control (*Luc* CDS) or bait (*Luc* CDS with *G3BP1* 5’ and 3’ UTRs) transcripts were pulled down from either WT or MAGE-B2 KO U2OS cells and subjected to immunoblotting for DDX5, MAGE-B2 or TRIM28 as a negative control. (E and F) DDX5 enhances G3BP1 translation *in vitro*. *In vitro* translation assays were performed using rabbit reticulocyte lysate with increasing amounts of recombinant GST (control), DDX5, or MAGE-B2 to measure translation of a luciferase reporter (n = 3) containing *G3BP1* 5’ and 3’ UTRs (E), or only *G3BP1* 5’ UTR (F). *P* values were determined by *t* test. (G and H) DDX5 helicase activity is required for enhancing G3BP1 translation *in vitro*. *In vitro* translation assays were performed using increasing amounts of recombinant GST (control), DDX5 WT, or DDX5 K144N (helicase dead mutant) to measure translation of a luciferase reporter (n = 3) containing *G3BP1* 5’ UTR (G) or *G3BP1* 5’ UTR lacking the DDX5 binding motif (10 nucleotide deletion) (H). *P* values were determined by *t* test. (I) *In vitro* translation assays reveal competition between DDX5 and MAGE-B2 for G3BP1 regulation via the 5’ UTR*. In vitro* translation assays were performed using recombinant DDX5 and titrating increasing amounts of GST (control) or MAGE-B2 to measure translation (n = 3) of a luciferase reporter containing *G3BP1* 5’ UTR. *P* values were determined by *t* test. (J) Regulation of G3BP1 translation via the 5’ UTR. Basal translation of the various luciferase reporters relative to control (Luc CDS) reveals that the presence of *G3BP1* 5’ UTR suppresses translation (n = 3). *P* values were determined by one-way ANOVA. (K) Model of the mechanism by which MAGE-B2 reduces G3BP and suppresses SG. MAGE-B2 inhibits G3BP translation by competing with the translational activator DDX5. This results in reduced G3BP protein levels, suppression of SG, and increased cellular stress threshold. Data are mean ± SD. Asterisks indicate significant differences from the control (* = p *≤* 0.05, ** = p *≤* 0.01, *** = p *≤* 0.001, n.s. = not significant) See also Figure S4.

To further characterize the mechanism by which MAGE-B2 and DDX5 regulate G3BP1 translation, we performed *in vitro* translation assays using a luciferase reporter in which the *G3BP1* 5’ UTR, 3’ UTR, both, or neither were included. We found that recombinant MAGE-B2 had no effect on translation of the reporters (Figure 4E). However, recombinant DDX5 promoted translation of the reporter (Figures 4E and 4F). Furthermore, this activity was dependent on the 5’ UTR of *G3BP1* (Figures 4F and S4A-B). In addition, the helicase activity of DDX5 was required for enhanced translation *in vitro*, as addition of DDX5 K144N had minimal effect on translation in comparison to wildtype DDX5 (Figure 4G). Previous reports have identified DDX5 binding motif consensus sequences (Lee et al., 2018). Scanning of the *G3BP1* 5’ UTR revealed a putative 10 nucleotide DDX5 recognition sequence (Figure 4H). Deletion of the putative DDX5-binding motif within the *G3BP1* 5’ UTR ablated DDX5’s ability to increase translation (Figure 4H).

Finally, we utilized the *in vitro* translation system to test for competition between MAGE-B2 and DDX5. Indeed, titrating increasing amounts of MAGE-B2 could inhibit DDX5’s ability to promote translation (Figures 4I and S4C-S4E). Consistent with a competition model, we did not observe binding between MAGE-B2 and DDX5 (Figures S4F-S4I). These results suggest that DDX5 is a key factor in determining G3BP1 translation and that this can be modulated by MAGE-B2 competition for *G3BP1* 5’ UTR binding. Interestingly, when comparing the basal translation of our reporter constructs, we found that the *G3BP1* 5’ UTR had a suppressive effect on translation (Figure 4J), which might suggest that the 5’ UTR contains a structural element that must be unwound by DDX5 for efficient translation. Overall, these findings suggest that MAGE-B2 and DDX5 act as key regulators of G3BP concentration to fine-tune the cellular stress response through controlling SG assembly dynamics (Figure 4K).

### The mouse ortholog of human MAGE-B2 (Mage-b4) regulates stemness of spermatogonial stem cells

Given our findings that MAGE-B2 regulates the SG response in cells, we sought to determine how this mechanism relates to normal physiology. To examine the expression pattern of *MAGE-B2*, we analyzed a panel of 26 human tissues by RT-qPCR and found that human *MAGE-B2* is restricted to expression in the testis (Figure 5A), which was consistent with *MAGE-B2* expression data from the Genome Tissue-Expression (GTEx) dataset of 53 disease-free human tissues (Figure S5A). We extended these analyses to characterize its expression profile in two mouse strains (BALB/C and C57BL/6) by RT-qPCR and found that the mouse ortholog of human *MAGE-B2*, *Mage-b4*, is predominantly expressed in the testis (Figures 5B and S5B). Notably, we found that *Mage-b4* underwent a gene duplication event in the mouse that resulted in a second copy (paralog), *Mage-b10*, which is 100% identical to *Mage-b4* within the MHD and only varies in the number of repetitive C-terminal elements that results in an 88 amino acid deletion (Figure S5C). Consistently, *Mage-b10* was also largely restricted to expression in the testis (Figure S5D). Given their high sequence identities and similar expression profiles, we refer to these two genes simply as *Mage-b4* herein.

**Figure 5.**
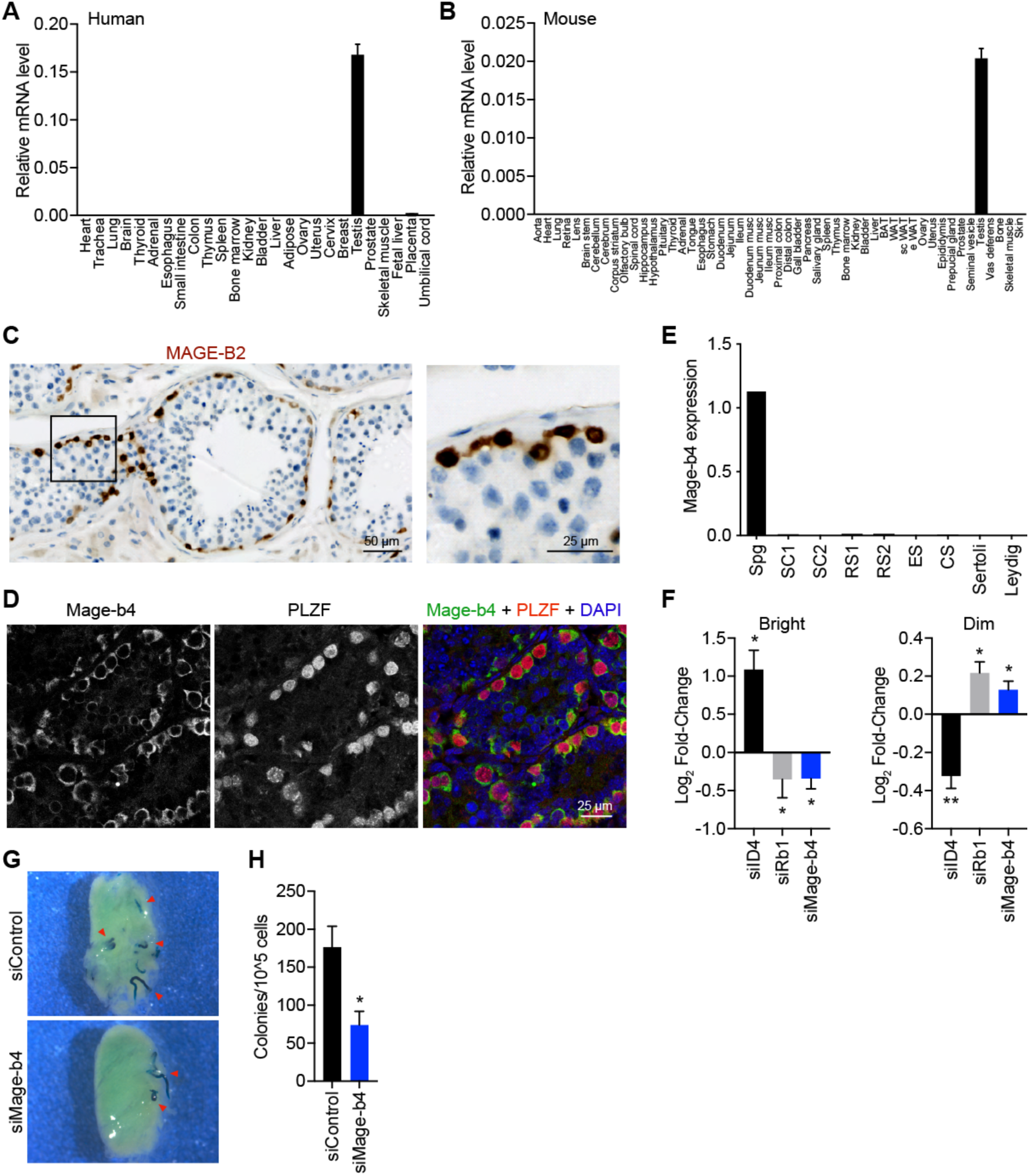
The mouse ortholog of human MAGE-B2 (Mage-b4) regulates stemness of testis spermatogonial stem cells. (A) Human *MAGE-B2* is expressed specifically in the testis. RT-qPCR analysis (n = 3) of normalized human *MAGE-B2* expression in the indicated tissues. Data are mean ± SD. (B) Mouse *Mage-b4* is expressed specifically in the testis. RT-qPCR analysis (n = 3) of normalized mouse *Mage-b4* expression in the indicated tissues from BALB/C mice. Data are mean ± SD. (C) Immunohistochemistry staining of human testis shows MAGE-B2 is expressed in spermatogonia. (D) Mouse Mage-b4 is expressed in undifferentiated spermatogonia. Mouse testis were immunostained for Mage-b4 and PLZF and representative images are shown. (E) Mouse Mage-b4 is enriched in spermatogonia. Analysis of previously described (Lukassen et al., 2018) single-cell RNA sequencing data derived from 8-week-old C57Bl/6J mice is shown. Spg = spermatogonia, SC = spermatocytes, RS = round spermatids, ES = elongating spermatids, CS = condensed/ condensing spermatids. (F) Mage-b4 maintains ID4-EGFP^bright^ stem cells in primary cultures of undifferentiated spermatogonia. Primary spermatogonia cultures were treated with the indicated siRNAs for 6 days before flow cytometry analysis to determine the percentage of ID4-EGFP^bright^ and ID4-EGFP^dim^ (n = 3). Log_2_ fold change of ID4-EGFP^bright^ (left) and ID4-EGFP^dim^ (right) is shown. Data are mean ± SEM. (G and H) Mage-b4 is required for efficient repopulation of testis. Spermatogonial transplantation assays were performed by transfecting LacZ-expressing primary spermatogonia cells with the indicated siRNAs for 6 days and transplanting them into the testes of recipient males depleted of germ cells. Two months later the number of LacZ-positive donor-derived colonies of spermatogenesis in the testis was determined (n = 3). Representative images are shown in (G) and quantification is shown in (H). Data are mean ± SEM. Asterisks indicate significant differences from the control (*p* values were determined by *t* test; * = p *≤* 0.05). See also Figure S5.

We next sought to determine in which cell type MAGE-B2 and its ortholog are expressed in the testis. Immunohistochemistry staining of human testis sections revealed that MAGE-B2 protein is enriched in spermatogonia (Figure 5C). Likewise, immunohistochemistry analysis of the mouse testis revealed that Mage-b4 co-localized with PLZF, a marker for undifferentiated spermatogonia including spermatogonial stem cells (SSC) (Osterlund et al., 2000) (Figure 5D). These findings are consistent with previous work from our lab demonstrating that Mage-b4 is enriched in spermatogonia based on developmental timing, cell sorting, and Kitl^Sl^/Kitl^Sl-d^ (steel) mice lacking spermatocytes (Fon Tacer et al., 2019). Furthermore, previously published single cell RNA-sequencing data also suggest that Mage-b4 is primarily expressed in spermatogonia (Figure 5E) (Lukassen et al., 2018). Additionally, single cell RNA-sequencing results identified Mage-b4, along with PLZF, as a unique marker of undifferentiated mouse spermatogonia (Jung et al., 2019). MAGE-B2 was also shown to be enriched in spermatogonia in human testis by single cell RNA-sequencing (Guo et al., 2018; Sohni et al., 2019; Xia et al., 2020).

To determine whether Mage-b4 expression is important for SSC maintenance, we utilized primary cultures of undifferentiated spermatogonia from mice expressing an Id4-eGFP reporter transgene (Chan et al., 2014). Knockdown of Mage-b4 decreased the fraction of EGFP^bright^ SSCs and increased the fraction of EGFP^Dim^ progenitor cells to a similar degree as a known regulator of SSC maintenance, Rb1 (Yang et al., 2013) (Figure 5F). Knockdown of Id4 was included as a positive control that increases the EGFP^Bright^ cell population due to a compensatory up-regulation of Id4-eGFP (Oatley et al., 2011). To determine whether Mage-b4 affects stem cell function *in vivo*, we performed spermatogonial transplantation assays (Brinster and Zimmermann, 1994) in which we depleted Mage-b4 in primary cultures of spermatogonia expressing a Rosa26-LacZ transgene before transplantation into the testes of recipient males lacking germ cells. The efficiency of the transplanted cells to repopulate the recipient males’ testes was determined by counting the number of LacZ-positive colonies clonally derived from a single transplanted SSC. Knockdown of Mage-b4 resulted in a significantly reduced ability to repopulate testes after transplantation into mice (Figure 5G and 5H), demonstrating a key role for Mage-b4 in regulating SSC function both *in vitro* and *in vivo*.

### The MAGE-B2 ortholog in mouse protects the male germline from heat stress

Given our results that MAGE-B2 regulates the stress response through repression of G3BP translation and SG, we reasoned that MAGE-B2 and its ortholog Mage-b4 may be important for controlling SG dynamics in SSC. Therefore, we utilized CRISPR-Cas9 to generate mice deficient in Mage-b4 and its paralog Mage-b10 (Figure S6A) and test their role *in vivo*. To confirm that Mage-b4 is the functional ortholog of human MAGE-B2, we treated primary cell cultures of undifferentiated spermatogonia from wildtype or Mage-b4 knockout mice with sodium arsenite and analyzed SG formation. Consistent with human MAGE-B2 function, we found that Mage-b4 knockout SSC exhibited increased SG formation (Figures 6A and S6B). Furthermore, we determined whether Mage-b4 regulates SG assembly in response to heat stress *in vivo*. Wildtype or Mage-b4 knockout mice were partially submerged in a control (33 °C) or heated water bath (38 °C or 42 °C) for 15 minutes to heat stress their lower extremities, including the testes. SG formation was analyzed by G3BP1 immunostaining of the testes. In line with the enhanced SG induction seen in Mage-b4 knockout primary cell culture, we found that the Mage-b4 knockout mice exhibited increased SG formation in response to heat, specifically in spermatogonia that normally express Mage-b4 (Figures 6B-6C and S6C).

**Figure 6.**
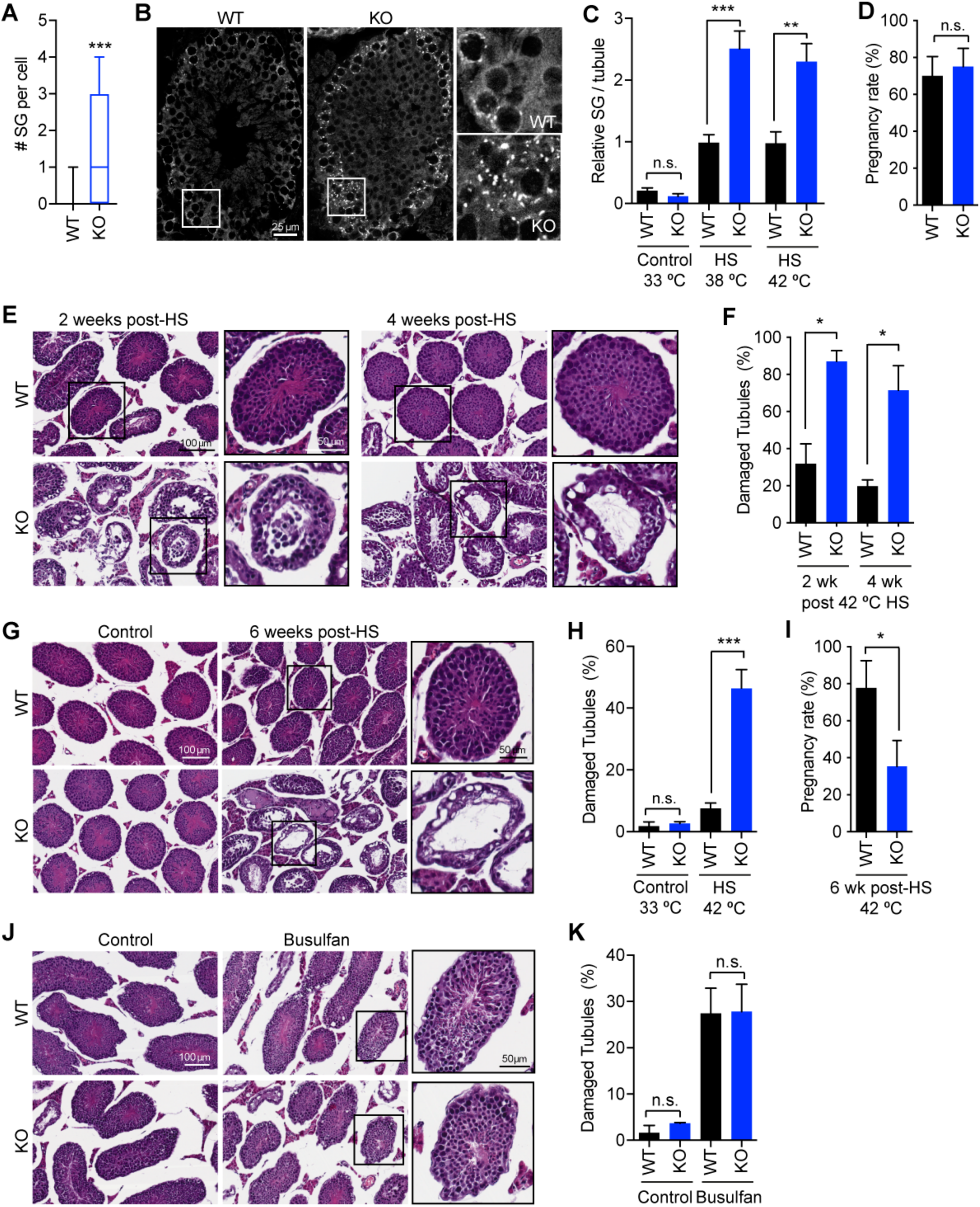
Knockout of the MAGE-B2 ortholog in mice results in hypersensitivity of the testis to heat stress. (A) Primary cultures of undifferentiated spermatogonia from WT or Mage-b4 KO mice were treated with 250 μM sodium arsenite for 1 hr and immunostained for PABP-C1. Quantification of SG number per cell from one representative experiment is shown. (B-C) Testes of WT or Mage-b4 KO mice were heat stressed (HS) for 15 min at 38 °C. Mice were immediately sacrificed, testes isolated and sectioned before G3BP1 immunostaining to detect stress granules. 10-20 seminiferous tubules were counted in testes of 4 mice per genotype totaling 80 tubules counted. Representative images (B) and quantitation (C) are shown. (D) Mage-b4 is not required for fertility under unstressed conditions, Pregnancy rates were determined by breeding WT or KO male mice to virgin WT females (20 mating trails across 12 mice per genotype). (E-H) Mage-b4 KO mice are hypersensitive to heat stress. WT or KO mice were control treated at 33 °C or heat shocked at 42 °C for 15 min and allowed to recover for 2, 4, or 6 weeks as indicated. Representative histology images (E, G) and quantification of damaged tubules (F, H) are shown (n = 6 control mice per genotype, n = 3-6 heat shocked mice per genotype per time point). (I) Mage-b4 KO mice are less fertile after heat stress. WT or KO mice were heat shocked at 42 °C for 15 min and allowed to recover for 6 weeks (n = 9 per genotype). P value was determined by Chi-square analysis. (J-K) Mage-b4 KO mice are not hypersensitive to genotoxic stress. WT or KO mice were intraperitoneally injected with 20 mg/kg body weight of busulfan and allowed to recover for 8 weeks before histological analysis of the testis. Representative images (J) and quantitation of damaged tubules (K) are shown (n = 2 control mice per genotype, n = 6 busulfan-treated mice per genotype). Data are mean ± SD. Asterisks indicate significant differences from the control (*p* values were determined by unpaired *t* test; * = p *≤* 0.05, *** = p *≤* 0.001, n.s. = not significant). See also Figure S6.

General characterization of the Mage-b4 knockout mouse revealed a modest reduction in testis weight (Figure S6D) whereas various other tissues were unaffected (Figures S6E-S6J). Fertility tests including pregnancy rate, litter size, sperm counts, and sperm motility revealed no significant defects in the fertility of male Mage-b4 knockout mice under standard laboratory conditions (Figures 6D and S6K-S6N). These data suggest that Mage-b4 is not required for male fertility under non-stressed conditions. This finding may not be surprising given the recent evolution of Mage-b4 in eutherian mammals (Katsura and Satta, 2011; Lee and Potts, 2017).

Spermatogenesis is a highly thermo-sensitive process such that maintenance of the testes at a temperature 4-5 °C lower than core body temperature is required for proper sperm production (Widlak and Vydra, 2017). Even modest increases in temperature can lead to apoptosis of germ cells, impaired spermatogenesis, and reduced testes weight (Reid et al., 1981; Rockett et al., 2001; Yin et al., 1997). Therefore, we speculated that Mage-b4 may have evolved in mammals to protect the male germline from mild heat stresses. Given that MAGE-B2 inhibition of SG formation is important for protecting cells against prolonged stress (Figures 1L and 2G), we determined whether Mage-b4 is important for spermatogenesis after heat stress. The testis of wildtype and Mage-b4 knockout mice were heat stressed for 15 min at 42 °C and were analyzed 2, 4, or 6 weeks later to allow enough time for two rounds of spermatogenesis. Strikingly, heat stress caused significantly greater reduction in the testis weight of Mage-b4 knockout mice compared to wildtype mice (Figure S6O). Histological analysis of testis from heat stressed mice revealed an increased number of damaged seminiferous tubules in Mage-b4 knockout mice compared to wildtype mice (Figures 6E-6H). Moreover, Mage-b4 knockout mice had reduced fertility after heat stress compared to wildtype mice (Figure 6I). Interestingly, Mage-b4 knockout mice treated with the genotoxic agent busulfan did not significantly differ from wildtype mice in terms of testis weight (Figure S6Q) or testis histology (Figures 6J and 6K), suggesting that SSC in Mage-b4 knockout mice are not generally more sensitive to all types of stress. In combination, these data suggest that Mage-b4 evolved to maintain the highly sensitive process of spermatogenesis in the face of fluctuating environmental temperatures by protecting germline cells in the testis.

## DISCUSSION

In this study, we identify a testis-specific factor, MAGE-B2, as a previously unknown regulator of the cellular stress response. Unlike previously characterized MAGE proteins, MAGE-B2 is an RBP that functions as a translational repressor of the essential SG nucleator, G3BP. Mechanistically, MAGE-B2 inhibits G3BP translation by competing with the translational activator, DDX5. Importantly, the fine-tuned suppression of SG that results from reduced G3BP protein levels increases the cellular stress threshold such that expression of MAGE-B2 allows for protection of the male germline from heat stress.

SG assembly is a multi-step process that typically begins with phosphorylation of the eukaryotic initiation factor 2α by a stress-sensing kinase to inhibit global translation (Sonenberg and Hinnebusch, 2009). The reduction of available 48S pre-initiation complex leads to polysome disassembly and the accumulation of untranslating mRNAs in the cytoplasm that can associate with G3BP and coalesce into SG cores (Kedersha and Anderson, 2002). These SG cores can then recruit additional IDR-containing RBPs, which mediate the assembly of additional proteins to form the mature SG consisting of a core and shell (Jain et al., 2016; Wheeler et al., 2016). As a highly coordinated biological process, SG can be regulated at multiple stages. Although knowledge of the biophysical drivers underlying SG formation (Kato et al., 2012; Molliex et al., 2015; Protter and Parker, 2016) and the RNA and protein composition of assembled SG is rapidly growing (Banani et al., 2016; Khong et al., 2017; Markmiller et al., 2018; Namkoong et al., 2018), our understanding of their regulation in typical physiological conditions is relatively limited. We find that MAGE-B2 modulates the stress response by directly regulating G3BP protein levels to suppress SG formation, thereby enhancing the tolerance to stress.

Like other RNP granules, SG are thought to form through the biophysical process of LLPS (Alberti et al., 2017; Protter and Parker, 2016). Phase separating proteins coalesce into liquid droplets *in vitro* when their concentration exceeds a critical threshold (Alberti, 2017). We demonstrate that this holds true for the key driver of SG phase separation, G3BP1. Moreover, we extend these *in vitro* studies to demonstrate the switch-like behavior between G3BP1 protein levels and SG initiation in cells and find that although MAGE-B2 has a modest effect on G3BP protein levels (two-fold), this change can have significant impact on SG formation.

What is the result of suppressed SG formation? Evidence supporting both pro-survival and pro-death functions of SG have been reported (Reineke and Neilson, 2019). However, a growing body of evidence suggests that disturbances in SG dynamics drives several age-related neurodegenerative diseases including amyotrophic lateral sclerosis, frontotemporal dementia, and inclusion body myopathy (Hackman et al., 2013; Mackenzie et al., 2017; Patel et al., 2015; Ramaswami et al., 2013). A number of disease-causing mutations affect SG disassembly and clearance and have been suggested to evolve into the aggregate pathology due to the increased risk of uncontrolled protein aggregation that accompanies high local concentrations of intrinsically disordered proteins. While it is not clear how the formation of these stable aggregates leads to the associated diseases, one model proposes that the increased predisposition to aggregate promotes promiscuous interactions within SG and increases the likelihood of SG to accumulate misfolded proteins, thereby impairing SG function. Likewise, disrupted SG clearance could compromise the function of RBPs that become trapped within aggregating SG and inhibit the synthesis of proteins essential for cell adaptation. Therefore, the suppression of SG and enhanced SG disassembly via regulation of G3BP protein levels by MAGE-B2 could reduce the risk of pathological aggregation, especially under chronic or repeated stress exposures that is common for the unique environment of SSC in the testis.

Recent biochemical and biophysical studies have demonstrated that MAGEs assemble with E3 RING ubiquitin ligases to form MAGE-RING ligases (MRLs) and function as regulators of ubiquitination in a multitude of cellular processes (Lee and Potts, 2017). The finding that MAGE-B2 regulates G3BP protein levels through translational regulation rather than ubiquitination was unprecedented. Approximately two thirds of MAGE-B2’s protein sequence consists of the highly conserved MHD that is shared among all human MAGEs. While it is possible that MAGE-B2’s MHD confers this unexpected function, it is likely that the N-terminal region outside the MHD contains an RNA recognition motif. In fact, the N-terminal region of MAGE-B2 (and mouse Mage-b4) includes a conserved 20 amino acid region enriched with basic residues (pI = 12.5), suggesting a potential interface for RNA binding. Whether other MAGEs, particularly those in the MAGE-B subfamily, share the capacity to bind RNA and the identification of other potential targets for translational repression will require further investigation.

DDX5, like other members of the DEAD box family of RNA helicases, is a multifunctional protein with roles in transcription as well as RNA and miRNA processing. Consistent with its functions in mediating various steps of gene expression, our results ascribe DDX5 with a role in translational regulation. *In vitro* translation assays revealed that DDX5 helicase activity was required for the enhanced translation of G3BP1 (Figure 4G). One possibility is that DDX5 unwinds and remodels a structural element within *G3BP1* 5’ UTR to allow for efficient translation. Interestingly, DDX5 was recently implicated as an inhibitor of translation (Hoch-Kraft et al., 2018). Further interrogation into how DDX5 achieves these opposing functions and the factors that dictate whether DDX5 activates or represses translation will provide informative insights into DDX5 mechanism of action.

Spermatogenesis is an intricate developmental process that depends on proper SSC maintenance to allow for the continuous production of sperm (de Rooij, 2017). In most mammals, testes are located in the scrotum outside the body, which allows spermatogenesis to occur at optimal temperatures substantially lower (4-5 °C) than the core body temperature (Widlak and Vydra, 2017). Spermatogenesis is highly thermo-sensitive such that elevated testicular temperature results in germ cell apoptosis, compromised sperm quality, and increased risk for infertility (Reid et al., 1981; Rockett et al., 2001; Yin et al., 1997). Although the detrimental effects of increased testicular temperature on spermatogenesis in mammals has been established for many years, the reasons why most mammals have evolved to maintain their testes at low temperatures remain unclear. Moreover, little is known about molecular mechanisms that have evolved to protect spermatogenesis from unstable temperature fluctuations, but regulation of SGs and mRNPs is an emerging concept for controlling germline stem cell homeostasis (Zhou et al., 2017). Given our findings that MAGE-B2-mediated modulation of G3BP and the stress response allowed for enhanced survival of cultured U2OS cells in the presence of oxidative stress (Figures 1L and 2G), we postulate that within the testis, SSC endogenously expressing MAGE-B2 utilize this function to allow for preservation of spermatogenesis in the presence of heat stress. Unfortunately, due to the embryonic lethality of G3BP1 knockout mice (Martin et al., 2013), precise determination of whether Mage-b4 protects germline cells through modulation of G3BP will not be trivial. Nonetheless, these findings suggest possible new routes for development of male fertility therapies through modulation of MAGE-B2, G3BP, and SG.

MAGEs are an evolutionarily ancient protein family, however, the type I MAGE CTAs only recently evolved in eutherians through a rapid expansion. Recent work from our lab revealed that the MAGE-A subfamily of MAGE CTAs evolved to protect primary spermatocytes against nutrient and genotoxic stress (Fon Tacer et al., 2019). Here, we propose that MAGE-B2 protects spermatogonial cells against heat stress through modulation of SG dynamics. Given the recent evolution of MAGE CTAs, it is perhaps not surprising that MAGE-B2 does not appear to impact basal spermatogenesis. Rather, MAGE-B2 serves as a mechanism to fine-tune the heat stress response in SSCs. This raises the question of whether such a mechanism might allow for the transmission of potentially damaged genetic material. Interestingly, it has been reported that heat stress does not induce pro-survival pathways via activation of heat shock transcription factors (HSFs) in meiotic and post-meiotic cells germ cells; rather, these cells undergo apoptosis, presumably to mitigate such unfavorable outcomes (Kus-Liskiewicz et al., 2013; Widlak and Vydra, 2017). Therefore, it is conceivable that the elevated heat stress threshold of SSC allows these select cells to survive and restore spermatogenesis, thereby preventing permanent azoospermia. Although most models of cellular stress management view each cell as autonomous sensory units, it is likely that this coordinated response provides a more systemic survival program to preserve fertility of the organism as a whole.

In summary, our work demonstrates that MAGE-B2 attenuates SG formation through translational inhibition of the SG nucleator, G3BP. Moreover, we provide evidence that the selective expression of MAGE-B2 in testis provides germline cells with an enhanced stress tolerance to maintain fertility in the face of stressful heat conditions, suggesting that the MAGE family evolved specifically to protect the male germline in eutherian mammals during times of stress.

## EXPERIMENTAL PROCEDURES

### Cell culture and transfection

U2OS, HCT116, HeLa, and HEK293 cells were grown in DMEM supplemented with 10% fetal bovine serum (FBS), 2 mM L-glutamine, 100 units/mL penicillin, 100 mg/mL streptomycin, and 0.25 mg/mL Amphotericin B. G3BP KO (both G3BP1 and G3BP2) U2OS cells and U2OS cells stably expressing G3BP1-GFP have been previously described (Figley et al., 2014; Zhang et al., 2018). siRNA and plasmid transfections were performed using Lipofectamine RNAiMAX (Invitrogen) and Effectene (QIAGEN), respectively, according to the manufacturers’ instructions.

### siRNAs

siRNAs were purchased from Sigma. Sequences of siRNAs: control: Sigma MISSION siRNA Universal Negative Control #1, MAGE-B2 #1: 5’-CGUUACAAAGGGAGAAAUG-3’, MAGE-B2 #2: 5’-GGCAGAUUCUUUACUUUGU-3’, MAGE-B2 #3: 5’-GUGGUCAAUUCUUGGUUUA-3’, G3BP1 #1: 5’-CAAAUCAGAGCUUAAAGAU-3’, G3BP1 #2: 5’-CUGAUGAUUCUGGAACUUU-3’, G3BP2 #1: 5’-CUCUGAACCAGUUCAGAGA-3’, G3BP2 #2: 5’-GAGCUAAAGGAAUUCUUCA-3’, DDX5 #1: 5’-CUGAUAGGCAAACUCUAAU-3’, DDX5 #2: 5’-CUACCUUGUCCUUGAUGAA-3’, DDX17 #1: 5’-GCCAAUACACCUAUGGUCA-3’, DDX17 #2: 5’-CCAAUACACCUAUGGUCAA-3’, DHX30 #1: 5’-GACAUCUUGCCCUUGGGCA-3’, DHX30 #2: 5’-CCUAUCACAGGCAAGCCCU-3’, HNRNPC #1: 5’-GAUGAAGAAUGAUAAGUCA-3’, HNRNPC #2: 5’-CUCAUUUAGUUGAGUAGCU-3’, SAFB #1: 5’-GAUGAUAAAUGUGACAGAA-3’, SAFB #2: 5’-GUAAUCCUGACGAAAUUGA-3’, ID4 #1: 5’-CAACAAGAAAGUCAGCAAA-3’, ID4 #2: 5’-GAGAUCCUGCAGCACGUUA-3’, ID4 #3: 5’-GCGAUAUGAACGACUGCUA-3’, ID4 #4: 5’-CCGUGAACAAGCAGGGUGA-3’, RB1 #1: 5’-GGAGUUUGAUUCCAUUAUA-3’, RB1 #2: 5’-GCAUAUCUCCGACUAAAUA-3’, RB1 #3: 5’-UCGAAGCCCUUACAAGUUU-3’, RB1 #4: 5’-UGCGUUAUCUACUGAAAUA-3’, Mage-b4 #1: 5’-AAGGAAGACAGGAATGCTGATG-3’, Mage-b4 #2: 5’-AGTCACACTTGTGGACTCTTGCA-3’, Mage-b4 #3: 5’-TGAGAATCCACAGAATGATCTT-3’, YB1 #1: 5’-CCUAUGGGCGUCGACCACA-3’, YB1 #2: 5’-GUUCCAGUUCAAGGCAGUA-3’, YB1 #3: 5’-GAAGUACCUUCGCAGUGUA-3’, AGO2 #1: 5’-GGUCUAAAGGUGGAGAUAA-3’, AGO2 #2: 5’-GGAUUCACGAGACCAGCUA-3’, and AGO2 #3: 5’-CCAUGUUCCGGCACCUGAA-3’.

### Antibodies

Rabbit polyclonal anti-MAGE-B2 antibody was produced by immunization of rabbits (Cocalico Biologicals) with recombinant full-length human MAGE-B2 protein produced in bacteria (described below) and affinity purified from serum. Antibody recognizing mouse Mage-b4 was previously described (Osterlund et al., 2000) and generously provided by Katarina Nordqvist. Commercial antibodies used in this study are as follows:

anti-Actin (Abcam, ab6276), anti-CAPRIN1 (Proteintech, 15512-1-AP), anti-DDX3 (Abcam, ab128206), anti-DDX5 (Abcam, ab126730), anti-eIF2α (Cell Signaling Technology, 5324), anti-eiF4G (Santa Cruz Biotechnology, sc-11373), anti-FLAG (Sigma, F3165), anti-G3BP1 (Abcam, ab181149), anti-G3BP1 (BD Biosciences, 611126), anti-G3BP1 (mouse tissue, BioRad VPA00492), anti-G3BP2 (Bethyl Laboratories, A302-040A), anti-GST (Potts and Yu, 2005), anti-HA (Sigma, A2095), anti-Myc (Santa Cruz Biotechnology, sc-40), anti-PABP-C1 (Abcam, ab21060), anti-PLZF (R&D Systems AF2944), anti-TIA1 (Santa Cruz Biotechnology, sc-166247), anti-TIAR (BD Biosciences, 610352), anti-TRIM25 (Abcam, ab167154), anti-TRIM28 (Abcam, ab22553), anti-Tubulin (Sigma, T9026), anti-USP10 (Proteintech, 19374-1-AP), anti-YB1 (Abcam, ab12148), anti-YTHDF1 (Proteintech, 17479-1-AP), anti-YTHDF2 (Proteintech, 24744-1-AP), anti-YTHDF3 (Santa Cruz Biotechnology, sc-377119), normal mouse IgG (Santa Cruz Biotechnology, sc-2025), and normal rabbit IgG (Santa Cruz, sc-2027).

### Generation of MAGE-B2 KO cells

MAGE-B2 knockout U2OS cells were generated using CRISPR-Cas9 technology. Briefly, two sgRNAs were used to delete the entire ORF (upstream sgRNA – 5’-GAAUAGAUGGUUAGUAUACC-3’, downstream sgRNA – 5’-UUUGGGAGAUUGAUUGGCUA-3’). 400,000 cells were transiently co-transfected with 200 ng of each gRNA expression plasmid (cloned into Addgene plasmid #43860), 500 ng Cas9 expression plasmid (Addgene plasmid #43945), and 200 ng of pMaxGFP via nucleofection (Lonza, 4D-Nucleofector X-unit) using solution P3, program CM-104 in small cuvettes according to the manufacturer’s recommended protocol. Five days post-nucleofection, cells were single-cell sorted by FACs for transfected cells based on pMaxGFP expression into 96-well plates. After sorting, cells were clonally expanded and screened for the desired modification using PCR-based assays. Knockout of MAGE-B2 was further verified by immunoblotting. Knockout lines with deletions were established. The sequences are indicated below: Clone #1: 5’–TCACAGATCTCATTCTCCCATCTCCAGGTA---deletion---ATCAATCTCCCAAAGCCAAGTTTACCTGCTGTT-3’ Clone #2: 5’–TCACAGATCTCATTCTCCCATCTCCA-----deletion--------ATCAATCTCCCAAAGCCAAGTTTACCTGCTGTT-3’.

### Immunostaining and stress granule analysis

Cells were washed with PBS, fixed in methanol for 10 min at −20 °C, permeabilized with blocking solution (PBS containing 0.2% (v/v) Triton X-100 and 3% (w/v) bovine serum albumin (BSA)) for 20 min at 4 °C, and incubated overnight at 4 °C with primary antibodies. The next day, cells were incubated with secondary antibodies conjugated with Alexa Fluor 488 or 568 for 30 min and nuclei were stained with DAPI. Stained cells were then imaged using a Leica SP8 TCS STED 3X confocal microscope. Stress granules were manually quantified using the indicated markers. At least 50-100 cells were counted for each condition in each experiment. For testis tissue sections, stress granules were quantified per seminiferous tubule using automated software. G3BP1 puncta were identified and segmented using IgorPro software. Segmented images were then counted using CellProfilerPro and the number of G3BP1 puncta (stress granules) was normalized to tubule size. 10-20 tubules were counted per mouse testis with at least four mice per genotype analyzed.

### Live-cell imaging and analysis

Live-cell imaging experiments were performed using either a Marianas 2 spinning disk confocal microscope or Bruker Opterra II Swept Field confocal microscope. Images were acquired using a 63x/1.4 Plan Apochromat objective with Definite focus or 60x Plan Apo 1.4NA oil objective with Perfect focus, respectively. During imaging, cells were maintained at 37 °C and supplied with 5% CO_2_ with an environmental control chamber and imaged at 40 s intervals with a 100 ms exposure time. For experiments measuring stress granule assembly and disassembly in response to heat stress, an objective heater and temperature-controlled flow chamber (Bioptechs) was utilized. For experiments correlating G3BP1 expression to stress granule initiation time, G3BP KO U2OS cells were transfected with GFP-G3BP1. 48 hr post-transfection, cells were treated with either 62 μM or 500 μM sodium arsenite. GFP intensity at t = 0 was measured as a readout of GFP-G3BP1 expression.

To determine if the relative amounts of G3BP1 in either WT or MAGE-B2 KO U2OS cells, G3BP KO U2OS cells were transfected with GFP-G3BP1. The following day, either WT or MAGE-B2 KO U2OS cells were seeded with the GFP-G3BP1 transfected cells. 48 hr post-transfection, cells were imaged to measure GFP intensity as a readout of GFP-G3BP1 expression. Cells were subsequently fixed and stained with a G3BP1 antibody as described above. Each cell that was previously imaged for GFP, was then imaged to measure G3BP1 intensity as a readout of G3BP1 expression. Linear regression analysis was applied to model the relationship between GFP-G3BP1 intensity and G3BP1 intensity in the G3BP KO cells transfected with GFP-G3BP1 (R^2^= 0.86). The calculated linear regression line (y=0.5308x + 640.28) was then used to determine the GFP-G3BP1 equivalent of the average measured endogenous G3BP1 intensity in WT or MAGE-B2 KO cells (8.0 or 13.4 RFU, respectively).

### Tandem affinity purification and mass spectrometry

Tandem affinity purification (TAP) was performed as described in previously (Doyle et al., 2010). Briefly, 15 15 cm^2^ dishes of HEK293 stably expressing TAP-Vector or TAP-MAGE-B2 were lysed, bound to IgG sepharose beads (GE Amersham), cleaved off the beads with TEV protease, collected on calmodulin sepharose beads (GE Amersham), eluted with SDS sample buffer, separated by SDS-PAGE and stained with colloidal Coomassie blue (Pierce). Total protein bands were excised, in-gel proteolyzed, and identified by LC/MS-MS.

### Cell viability

Cells were seeded in 6-well plates and allowed to adhere for at least 4 hr before being treated with 4 μM sodium arsenite for 72 hr. Cells were trypsinized and the number of viable cells was determined using a Beckman Coulter Vi-Cell XR automated cell counting system.

### Preparation of cell lysates and immunoblotting

Cells were washed with PBS, collected by scraping, and pelleted by centrifugation. Cell pellets were lysed in NP-40 lysis buffer (50 mM Tris-HCl pH 7.5, 150 mM NaCl, 0.5% (v/v) IGEPAL CA-630 (Sigma-Aldrich), 1 mM DTT, and 1X protease inhibitor cocktail (Sigma-Aldrich)) for 30 min on ice and centrifuged to clarify. The Micro BCA Protein Assay Kit (Thermo Scientific) was used to quantify total protein concentration of lysates. Lysates were prepared in SDS sample buffer, resolved on SDS-PAGE gels, and transferred to nitrocellulose membranes. Membranes were blocked with 5% BSA in TBST (25 mM Tris pH 8.0, 2.7 mM KCl, 137 mM NaCl, 0.05% (v/v) Tween-20) and incubated with primary antibodies. After three washes with TBST, membranes were incubated with secondary antibodies, washed an additional three times, and detected via chemiluminescence or near-infrared fluorescence.

### Protein disorder prediction

Intrinsically disordered regions were calculated using IUPred2A global structure disorder prediction (long disorder, default option). G3BP1 protein sequence was used as the input and the IUPred server returned a disorder tendency score between 0 and 1 for each residue with higher values corresponding to a higher probability of disorder.

### Recombinant protein purification

GST-MAGE-B2, GST-DDX5 (wildtype or K144N), or GST alone were produced in BL21-CodonPlus (DE3)-RIPL cells by overnight induction at 16 °C with 0.5 mM Isopropyl b-D-1-thiogalactopyranoside (IPTG). GST-G3BP1 was produced in BL21-CodonPlus (DE3)-RIPL cells using the ZYM-5052 complex auto-inducing medium for induction by lactose (Studier, 2014).

Bacterial pellets were solubilized with 50 mM Tris-HCl pH 7.7, 150 mM KCl, 1 mM dithiothreitol (DTT), and 1X protease inhibitor cocktail (Sigma) for GST and GST-MAGE-B2; 50 mM Tris-HCl pH 7.4, 300 mM KCl, 10% (v/v) glycerol, 1 mM DTT, and 1X protease inhibitor cocktail for GST-DDX5; and 50 mM Tris-HCl pH 8.0, 300 mM NaCl, 10% (v/v) glycerol, 1 mM DTT, and 1X protease inhibitor cocktail for GST-G3BP1. GST-tagged proteins were purified from bacterial lysates with glutathione Sepharose (GE Amersham) and eluted with 10 mM glutathione. In some cases, the GST tag was cleaved by on column digest overnight at 4 °C with Precision Protease. Purified proteins were immediately used for all assays prior to freezing.

### *In vitro* phase separation assay

Samples were prepared by combining the indicated concentrations of purified recombinant G3BP1 and NaCl in a buffer containing 50 mM Tris-HCl pH 8.0 and 100 mg/mL Ficoll 400 (Sigma-Aldrich). Samples were sandwiched between a hydrophobic coverslip and microscope slide using an adhesive imaging spacer prior to imaging on a Lecia SP8 Widefield microscope.

### RT-qPCR

RT-qPCR analysis was performed as described previously (Pineda et al., 2015). RNA was extracted using either RNAStat60 (TelTest) or Trizol Reagent (Invitrogen) according to the manufacturers’ instructions, treated with DNase I (Roche), and converted to cDNA using the High Capacity cDNA Reverse Transcription kit (Life Technologies). cDNA was subjected to qPCR and gene expression was measured using SYBR Green (Applied Biosystems). Data were analyzed by ΔΔCt method normalizing to *18S* rRNA. qPCR primers used: *G3BP1* Forward: 5’-TGAGGTCTTTGGTGGGTTTG-3’, *G3BP1* Reverse: 5’-TGCTGTCTTTCTTCAGGTTCC-3’, *18S* rRNA Forward: 5’-ACCGCAGCTAGGAATAATGGA-3’, *18S* rRNA Reverse: 5’-GCCTCAGTTCCGAAAACCA-3’, *RPLP0* Forward: 5’-TCTACAACCCTGAAGTGCTTGAT-3’, *RPLP0* Reverse: 5’-CAATCTGCAGACAGACACTGG-3’, *Mage-b4* Forward: 5’-TGAGCAAGCACCCATTACTTTG-3’, and *Mage-b4* Reverse: 5’-TGACGGTTTACACATTTCTCTTTGT-3’.

### ^35^S metabolic labeling

^35^S labeling experiments were performed as described previously (Bonifacino, 2001). Briefly, U2OS Cells were washed with PBS and incubated with labeling media (methionine- and cystine-free DMEM supplemented with 10% dialyzed FBS, 2 mM L-glutamine, 100 units/mL penicillin, 100 mg/mL streptomycin, and 0.25 mg/mL Amphotericin B) for 30 min at 37 °C. Cells were then labeled with 0.1 mCi/mL EasyTag EXPRESS ^35^S Protein Labeling Mix (Perkin Elmer) diluted in labeling media. For G3BP1 half-life experiments, cells were pulse labeled for 1 h at 37 °C, chased with complete media (DMEM supplemented with 10% FBS, 2 mM L-glutamine, 100 units/mL penicillin, 100 mg/mL streptomycin, and 0.25 mg/mL Amphotericin B), and harvested at the indicated time points. For G3BP1 translation experiments, cells were labeled and harvested at the indicated time points. Cells were lysed in RIPA lysis buffer (50 mM Tris-HCl pH 7.5, 150 mM NaCl, 1% (v/v) IGEPAL CA-630, 0.5% (w/v) sodium deoxycholate, 0.5% (w/v) SDS, 1 mM DTT, and 1X protease inhibitor cocktail) for 30 min on ice and centrifuged to clarify. Supernatants were pre-cleared with control IgG for 1 hr at 4 °C and Protein A/G Plus-Agarose beads (Santa Cruz Biotechnology) for 30 min at 4 °C before incubation with primary antibody overnight at 4 °C. The next day, samples were incubated with beads for 4 hr at 4 °C, washed with RIPA lysis buffer, and eluted with SDS sample buffer. Samples were resolved on SDS-PAGE gels, and either transferred to nitrocellulose membranes for immunoblotting, or fixed and dried for phosphorimaging. For global translation experiments, cells were labeled and harvested at the indicated time points. 10 µL of the labeled cell suspension were added to 0.1 mL of BSA/NaN_3_ (1 mg/mL BSA containing 0.02% (w/v) NaN_3_) and precipitated by the addition of 1 mL 10% (w/v) trichloroacetic acid (TCA). Samples were vortexed and incubated for 30 min on ice before filtration onto glass microfiber disks. Disks were washed with 10% (w/v) TCA and ethanol. 10 µL of the labeled cell suspension was spotted onto another glass microfiber disk to measure the total amount of radiolabeled amino acid. Disks were transferred to vials with scintillation fluid for scintillation counting.

### Co-immunoprecipitation

Cells were washed with PBS, collected by scraping, and pelleted by centrifugation. Cell pellets were lysed in NP-40 lysis buffer (50 mM Tris-HCl pH 7.5, 150 mM NaCl, 0.5% (v/v) NP-40, 1 mM DTT, and 1X protease inhibitor cocktail (Sigma-Aldrich)) for 30 min on ice and centrifuged to clarify. Soluble lysates were incubated with myc-conjugated Protein A beads for 2 hr at 4 °C. Beads were washed with NP-40 lysis buffer, eluted with SDS sample buffer, and resolved on SDS-PAGE for subsequent immunoblotting.

### Polysome profiling

Polysomal profiling in U2OS WT and MAGE-B2 KO cells was done as described previously (Karamysheva et al., 2018) with minor modifications specific to cell culture. Cells were seeded in 20 mL of the DMEM medium (Sigma-Aldrich) supplied with 10% FBS and penicillin/streptomycin mixture (100 units and 100 µg/mL correspondingly) (Sigma-Aldrich) with initial cell count of 0.5×10^5^ cells/mL. Cells were grown at 37 ^°^C and 5% CO_2_ for 5 days. To arrest the ribosome on translated mRNAs cells were treated with 100 µg/mL cycloheximide (CHX) (Sigma-Aldrich) for 10 min at 37 ^°^C and 5% CO_2_. After incubation with CHX cells were washed twice with cold PBS (Sigma-Aldrich) supplied with 100 µg/mL CHX, and immediately lysed on ice with 500 µl of 20 mM HEPES-KOH (pH 7.4), 100 mM KCl, 5 mM MgCl_2_, 1 mM DTT, 0.5% NP-40, 1x protease inhibitor cocktail (EDTA-free), 100 µg/mL CHX, and 1 mg/mL heparin (Sigma-Aldrich). Cells were scraped from the plate, and concentrations of MgCl_2_ and NP-40 were adjusted for increased sample volume. The cell lysates were passed through a 22-gauge needle 6 times and then clarified by centrifugation at 11,200 × g and 4 ^°^C for 8 min. To evaluate the amount of starting material for polysome fractionation, absorbance at 260 nm was measured in cell lysates and adjusted to have equal input for all samples. 500 µl of cell lysate was used for fractionation. Linear sucrose gradients were prepared using Gradient Master 108 (BioComp) with 10% and 50% sucrose containing 20 mM HEPES-KOH (pH 7.4), 100 mM KCl, 10 mM MgCl_2_, 1 mM DTT, 0.5% NP-40, 1x protease inhibitor cocktail (EDTA-free), and 1 mg/mL heparin in Polyclear centrifuge tube (Seton). Lysates were loaded on the top of the gradient and centrifuged at 260,000 × g and 4 ^°^C for 2 h using SW 41 rotor (Beckman). Collection of polysomal fractions were done using Piston Gradient Fractionator (BioComp). Absorbance at 260 nm and 280 nm was recorded during each run of fractionation.

### Affinity pulldown of biotinylated RNA

Affinity pulldown of biotinylated RNA for detection of protein-RNA complexes was performed as previously described (Panda et al., 2016). Briefly, pCS2-luciferase CDS or pCS2-luciferase with G3BP1 5’UTR and 3’UTR were linearized with Sal I (New England Biolabs) and gel purified (Qiagen). Biotinylated transcripts were produced from the linearized DNA *in vitro* using the MEGAscript kit (Life Technologies) and biotin-UTP (Sigma) according to the manufacturer’s instructions and purified by the MEGAclear spin columns (Life Technologies) according to the manufacturer’s instructions. Integrity of the transcripts were confirmed by non-denaturing agarose gel electrophoresis.

The purified control Luciferase CDS RNA or Luciferase CDS with *G3BP1* UTR RNA (10 µg) were added to cell lysate supernatants (approximately 1 mg) prepared from U2OS parental or MAGE-B2 KO cells in NP-40 lysis buffer (50 mM Tris-HCl pH 7.5, 150 mM KCl, 0.5% (v/v) NP-40, 1 mM DTT, 1X protease inhibitor cocktail (Sigma), and Ribolock RNase inhibitor (Thermo Fisher)). RNA was incubated in cell lysates for 45 min at room temperature followed by addition of 50 µL streptavidin Dynabeads M-280 (Invitrogen) for 45 additional minutes at room temperature. Beads were separated on a magnetic stand and washed 4 times with NP-40 lysis buffer. Proteins were eluted in 1X SDS-sample buffer and analyzed by liquid chromatography with tandem mass spectrometry (LC-MS/MS) or immunoblotting using the following antibodies: anti-DDX5 (Abcam, ab126730), anti-MAGE-B2 (described above), and anti-TRIM28 (Abcam, ab22553).

### CLIP-qPCR

Cross-linking immunoprecipitation and QPCR (CLIP-QPCR) was carried out as previously described (Yoon and Gorospe, 2016). Briefly, HEK293/TAP-Vector, HEK293/TAP-MAGE-B2, U2OS parental, or U2OS MAGE-B2 KO cells (10-15 15 cm^2^ dishes) were washed in ice-cold, magnesium-free PBS and irradiated on ice with 150 mJ/cm^2^ of UVC (254 nm) in a Stratalinker 2400 (Agilent). Cells were collected in ice-cold PBS, pelleted, lysed in NP-40 lysis buffer (50 mM Tris-HCl pH 7.5, 150 mM KCl, 0.5% (v/v) NP-40), and centrifuged for 15 min at 10,000 xg at 4 °C. Supernatants were collected and subjected to immunoprecipitation. HEK293 cell lysates were incubated with 20 µL IgG Sepharose (GE Healthcare) or as a control Sepharose without IgG (GE Healthcare) for 3 hr at 4 °C rotating. U2OS cell lysates were incubated with 20 µL pre-coupled antibody-protein A/G PLUS-agarose beads (Santa Cruz Biotechnology) for 3 hr at 4 °C rotating. Antibodies (10 µg) used were as follows: normal rabbit IgG control (Santa Cruz, sc-2027) and anti-DDX5 (Abcam, ab126730). Beads were then washed three times in NP-40 lysis buffer, treated with 20 units of RNase-free DNase I for 15 min at 37 °C, and proteins degraded by treatment with 0.5 mg/mL proteinase K (Invitrogen) in 0.5% SDS at 55 °C for 15 min. RNA was then separated by phenol:chloroform extraction, followed by ethanol precipitation. RNA was then converted to cDNA using the High Capacity cDNA reverse transcriptase kit (Invitrogen) according to manufacturer’s instructions. qPCR analysis was performed on cDNA using PowerUp SYBR Green master mix (Applied Biosystems) according to manufacturer’s instructions using the following primers: *G3BP1* F 5’-TGAGGTCTTTGGTGGGTTTG-3’, *G3BP1* R 5’-TGCTGTCTTTCTTCAGGTTCC-3’, *RPLP0* F 5’-TCTACAACCCTGAAGTGCTTGAT-3’, *RPLP0* R 5’-CAATCTGCAGACAGACACTGG-3’. Data were analyzed by ΔΔCt method normalizing to *RPLP0* and control pulldowns (normal IgG (U2OS) or non-IgG beads (HEK293)).

### *In vitro* translation assays

*In vitro* translation assays monitoring luciferase enzyme production in rabbit reticulocyte lysate were carried out as follows. Purified proteins, GST, GST-MAGE-B2, DDX5 WT, or DDX5 K144N, were added alone or in combination at the indicated concentrations (0.25-4 pmol) to *in vitro* translation reactions (Promega SP6-TNT Quick rabbit reticulocyte lysate system) containing firefly luciferase coding sequence (CDS) alone or luciferase CDS with *G3BP1* 5’ UTR and/or *G3BP1* 3’ UTR sequences. Reactions were incubated at 30 °C for 1.5 hr. Samples were diluted in passive lysis buffer (Promega) before analysis of luciferase protein levels using Dual-Glo luciferase assay system (Promega) and an EnSpire multimode plate reader (Perkin-Elmer).

### *In vitro* binding assays

*In vitro* binding assays were performed as described previously (Doyle et al., 2010; Hao et al., 2013). 15 μg of purified GST-tagged proteins were bound to glutathione Sepharose beads in binding buffer (25 mM Tris pH 8.0, 2.7 mM KCl, 137 mM NaCl, 0.05% (v/v) Tween-20, and 10 mM 2-mercaptoethanol) for 1 hr and then blocked for 1 hr in binding buffer containing 5% (w/v) milk powder. *In vitro* translated proteins (Promega SP6-TNT Quick rabbit reticulocyte lysate system) were then incubated with the bound beads for 1 hr, extensively washed in binding buffer, eluted with 2X SDS-sample buffer, boiled, subjected to SDS-PAGE, and immunoblotting.

### Animals and tissue collection

Human tissues were obtained from commercially available sources and mouse tissues were collected as described previously (Fon Tacer et al., 2019). All procedures and use of mice were approved by the Institutional Animal Care and Use Committee of St. Jude Children’s Research Hospital.

### Spermatogonial cultures, ID4-eGFP flow cytometry and transplantation analyses

Spermatogonia cultures were established as previously described (Fon Tacer et al., 2019). Briefly, primary cultures of undifferentiated spermatogonia were generated from the double-transgenic *Id4-eGFP;Rosa26-LacZ* hybrid mice that express the LacZ transgene in all germ cells, but express the EGFP transgene only in ID4+ spermatogonial stem cells (Chan et al., 2014; Helsel et al., 2017b) or Mage-b4 knockout mice (described below). Primary cultures were established from the magnetic-activated cell sorting (MACS)-sorted THY1+ fraction of testis homogenates of P6–8 mice (Oatley and Brinster, 2006). Spermatogonial cultures were maintained on mitotically inactivated SIM mouse embryo-derived thioguanine- and ouabain-resistant feeder monolayers (STOs) in mouse serum-free medium (mSFM), devoid of fatty acids and supplemented with the growth factors GDNF (20 ng/mL; Peprotech, NJ, USA) and FGF2 (1 ng/mL; Peprotech). Cultures were kept in glycolysis-optimized conditions in humidified incubators at 37 °C, 10% O_2_ and 5% CO_2_ in air (Helsel et al., 2017a). Culture media was replaced every other day, and passaging onto fresh feeders was performed every 6–8 days.

For siRNA transfection, cultured spermatogonial clumps were separated from feeders by gentle pipetting and a single-cell suspension was generated by trypsin-EDTA digestion. 1×10^5^ or 2×10^4^ cells were plated in 24 or 96 well plates, respectively, without STO feeder cells in mSFM with GDNF and FGF2. Cells were transfected with either non-targeting control (Dharmacon, D-001810-10-05) or Mage-b4/10 siRNA oligonucleotides by Lipofectamine 3000 (Invitrogen) as follows: 2 µL Lipofectamine 3000 in 100 µL OptiMem and 75 pmol siRNA in 100 µL OptiMem per 1×10^5^ cells. 16 hrs after transfection, cells were washed with HBSS and fresh mSFM with GDNF and FGF2 was added. To assess ID4-eGFP expression level, cells were analyzed by flow cytometry 6 days after transfection. Single-cell suspensions were generated by trypsin-EDTA digestion as described previously and analyzed using an Attune NxT Flow Cytometer (Thermo Scientific, MA, USA). Identification and gating of the ID4-eGFPbright and ID4-eGFPdim populations from cultures was done as described previously (Chan et al., 2014; Helsel et al., 2017b).

To compare the regenerative capacity of spermatogonial culture populations after siRNA transfection, transplantation analysis was performed (Oatley and Brinster, 2006). Six days after siRNA transfection spermatogonial clumps were separated from feeders by gentle pipetting, single-cell suspension generated by trypsin-EDTA digestion and cells suspended in mouse serum-free medium at 1×10^6^ cells/ml. 10 µl (10,000 cells) was microinjected into each recipient testis. Recipient testes were evaluated for colonies of donor-derived spermatogenesis 2 months later.

### Generation and genotyping of *Mage-b4* knockout mouse models

Mouse lines were generated by injecting sgRNAs and Cas9 protein in the pronucleus and cytoplasm of C57BL/6 zygote. A single sgRNA targeting both *Mage-b4* and *Mage-b10* (TCCAGAATCCTGATTAGAGC) was prepared by *in vitro* trascription (MEGAshortscript T7 Kit, Ambion) and purified by MEGAclear™ Transcription Clean-Up Kit (Ambion). The progeny was screened for frameshift mutations by Cel-1 assay and Sanger sequencing. Animals were genotyped as previously described (Fon Tacer et al., 2019). Briefly, tail snips (1-2 mm) were collected at weaning (∼21 days old) and again when animals were euthanized for organ collection. Genomic DNA (gDNA) was prepared by incubating tails in 200 μl of 50 mM NaOH at 95 °C for 35 min, allowed to cool for a 3 min, and neutralized by addition of 20 μl 1 M Tris (pH 8.0). PCR was performed using KAPA2G Robust Hotstart PCR kit (KAPA Biosystems, #KK5518), following the manufacturer’s protocol. Buffer A, Enhancer, and 1 μl of gDNA was used. Primers used were as follows: *Mage-b4* forward – 5’-TTCCTCAATCCCGGTACAAG-3’, *Mage-b4* reverse – 5’-TGTGCACCTTCCCATCATAA-3’, *Mage-b10* forward – 5’-TACTAAGCTAGCTCTAGCGG-3’, *Mage-b10* reverse – 5’-AAAGTACCAGAGGTCCAAGGGAGGA-3’. The *Mage-b4* locus was genotyped by PCR alone.

The *Mage-b10* locus was amplified by PCR and then amplicons were digested with BslI enzyme (NEB) at 55°C for 1 hour in NEB Cutsmart buffer; KO animals displayed a unique banding pattern compared to WT.

### Tissue weights and fertility evaluation

To assess the effect of Mage-b4 depletion on organ weights, wildtype and KO mice were sacrificed and organ weights measured immediately after dissection. To test the fertility in males, wildtype females were paired with individually housed wildtype or KO males and checked for vaginal plugs for up to 4 days. Females were then removed, and males were allowed two days to recover before addition of a new female. This process was then repeated for a total of 3 females per male. Females that were successfully plugged were then monitored for pregnancy. We recorded successful pregnancies, number of pups born, and average pup weight at birth. Sperm was prepared from cauda epididymis. Cauda epididymal sperm were allowed to swim out and were incubated for 15 min at 37 °C in DPBS (Dulbecco’s’ PBS with 0.1% FBS, 10 mM HEPES, 10 mM sodium pyruvate, glucose (1 mg/mL), and penicillin/streptomycin (1 mg/mL)). Sperm number and motility were quantified using a computer-assisted semen analysis system (Hamilton Thorne Research) and by manual counting using a hemocytometer. We used 6 males per genotype for fertility and organ weight measurement.

### Heat stress and busulfan treatments

To evaluate spermatogenesis recovery after heat stress 10-12 week old mice were anesthetized and placed in a polystyrene float in a water bath so that their lower third was submerged. Mice were anesthetized by intraperitoneal injection of ketamine:xylazine (100 mg/kg:10 mg/kg body weight).

Water baths were maintained at 33 °C for control or 38 °C or 42 °C for heat stress. Animals were incubated for 15 minutes, removed, dried, and (when appropriate) monitored for recovery. For monitoring stress granules, mice were sacrificed immediately after heat stress and processed for G3BP1 immunostaining as described below. At sacrifice (immediately or 2, 4, or 6 week post-stress), testes were weighed and fixed in 4% PFA for 24 hr. For busulfan treatment, 4 month old wildtype and KO mice were intraperitoneally injected with 20 mg/kg body weight of busulfan (Sigma) dissolved in 1:1 volume ratio of DMSO and water. Eight weeks after treatment, mice were sacrificed and analyzed similarly as heat stressed mice above.

### Immunohistochemistry (IHC) and immunofluorescence (IF) tissue staining

Testes were fixed for 24 hr at 4 °C in 0.1 M sodium phosphate buffer, pH 7.2, containing 4% (w/v) paraformaldehyde (PFA). Fixed testes were paraffin embedded for H&E staining and IHC. IHC-based labeling was performed after de-paraffinization using a Discovery XT autostainer (Ventana Medical Systems) with anti-Mage-b4 antibody. All slides were counterstained with hematoxylin. Bright-field images were taken with an upright Eclipse Ni (Nikon) or constructed from digitized images using Aperio ImageScope (Leica Biosystems). The percentage of damaged tubules showing vacuolization and reduced germ cell layers (<4) was determined in a blinded manner in which >200 tubules from 2-6 mice per genotype per experiment were analyzed.

For IF, fixed testes were incubated in a 10% sucrose solution (w/v, dissolved in 1x PBS) at 4 °C until equilibrated, and then in 30% sucrose overnight at 4 °C. Once equilibrated, testes were embedded in tissue freezing medium (Electron Microscopy Sciences) and frozen using a Shandon Lipshaw cryobath. Tissues were cut into 6 µm sections. Prior to labeling, sections were equilibrated in air to room temperature for 8 min, hydrated for 8 min in PBS at room temperature, heat-treated at 80 °C for 8 min in 10 mM sodium citrate (pH 6.0) and then incubated for 1 hour at room temperature in blocking buffer (3% (w/v) BSA or 1% (v/v) Roche Blocking Reagent, diluted in 0.1 M sodium phosphate buffer, containing 0.2% (v/v) Triton X-100. Sections were then treated for 18-24 hr at 4 °C with antibody diluted in blocking buffer. Antibodies used were as follows: anti-Mage-b4 (Nordqvist lab, 1:100), anti-PLZF (R&D Systems AF2944, 1:50), and anti-G3BP1 (BioRad VPA00492, 1:100). After treatment with primary antibodies, sections were washed 3 times for 10 min per wash in PBS containing 0.02% (v/v) Triton X-100 and then incubated for 1 hr at room temperature with Alexa-flour secondary antibodies (Molecular Probes) diluted to 4 µg/mL in blocking buffer. After treatment with secondary antibodies, sections were washed two times in PBS containing 0.02% (v/v) Triton X-100. Samples were then incubated in DAPI diluted to 1 μg/ml in PBS for 5 minutes at room temperature, washed once in PBS containing 0.02% (v/v) Triton X-100, once in PBS, and cover-slipped for viewing using Fluorogel mounting medium (Electron Microscopy Sciences).

## QUANTIFICATION AND STATISTICAL ANALYSIS

All data were analyzed by Prism 7 (GraphPad). Statistical details of experiments can be found in figure legends. Unless noted otherwise, data are representative of at least three biologically independent experiments. Single time-point datasets were analyzed by *t* test. Multiple comparison (>3 groups) was performed using one-way Anova with Dunnett’s post-hoc test as indicated. Multiple time-point datasets were analyzed by two-way ANOVA unless otherwise stated. For all statistical analyses: * = p *≤* 0.05, ** = p *≤* 0.01, *** = p *≤* 0.001, **** = p *≤* 0.0001, n.s. = not significant.

## ACKNOWLEDGEMENTS

We thank members of the Potts lab for helpful discussions and critical reading of the manuscript. We would also like to thank Dr. Katarina Nordqvist (Nobel Museum, Sweden) for generously providing Mage-b4 antibody. We thank the St. Jude Proteomics Facility for the mass spectrometry-based protein analysis and the Transgenic/Gene Knockout Shared Resource for generation of Mage-b4 knockout mice.

## Funding

This work was partially supported by American Cancer Society Research Scholar Award 181691010 (P.R.P.).

## Author contributions

P.R.P. and A.K.L. conceptualized the study and designed experiments. S.N.P. and S.M.P.-M. performed gene editing. K.F.T., T.L., M.J.O., and J.M.O. performed SSC and transplantation assays. J.K. and K.F.T. developed the mouse model and performed *in vivo* analysis. E.B.T. and A.L.K. performed polysome analysis. P.Y., H.J.K., and J.P.T. provided critical reagents and project guidance. A.K.L. and P.R.P. performed cell culture and in vitro experiments. A.K.L. and P.R.P. wrote the manuscript.

## Competing interests

The authors have no competing interests.

