## Supplemental Figures for "Enhanced stress tolerance through reduction of G3BP and suppression of stress granules"

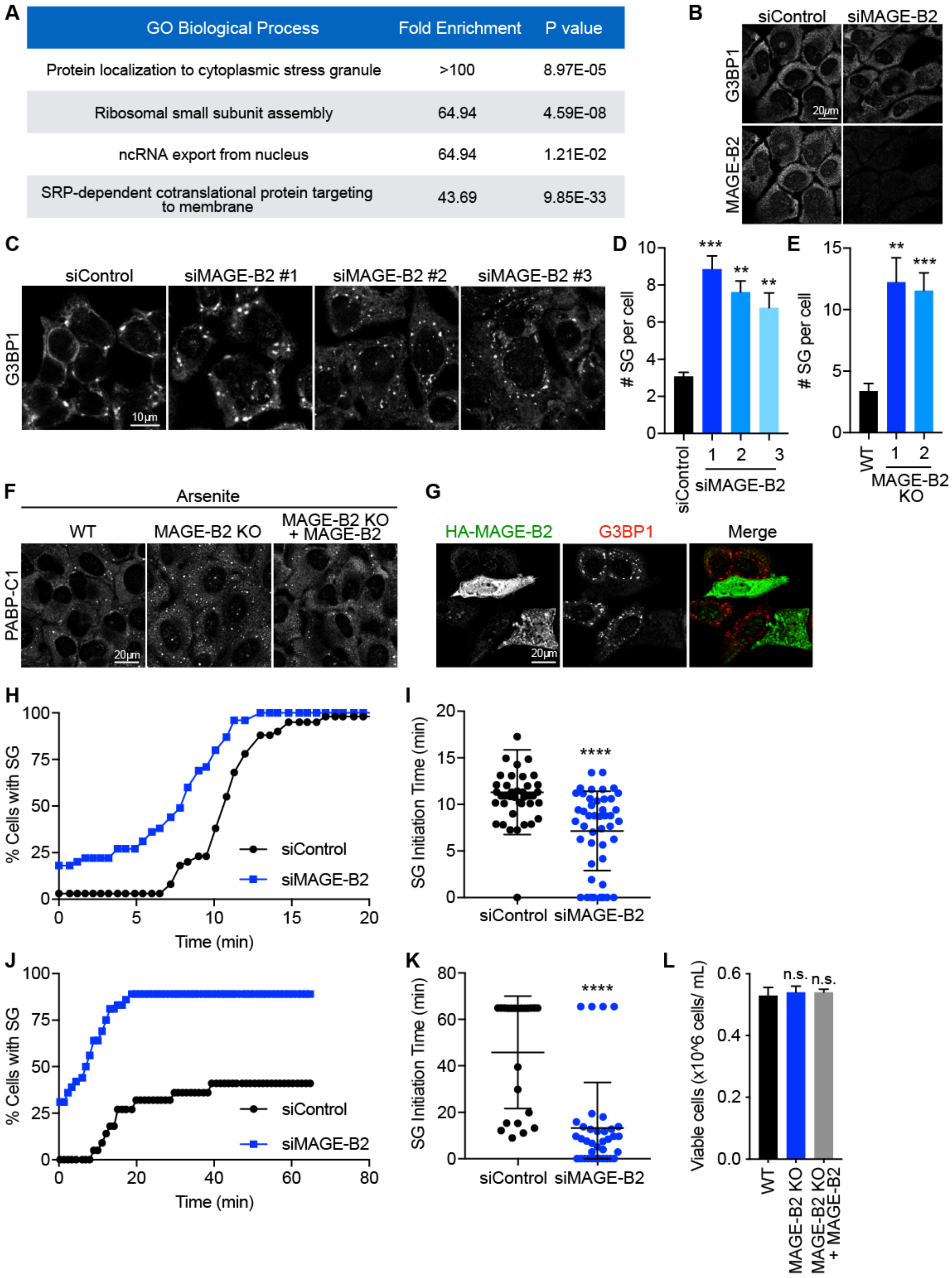

| GO Biological Process | Fold Enrichment | P value |
|---|---|---|
| Protein localization to cytoplasmic stress granule | >100 | 8.97E-05 |
| Ribosomal small subunit assembly | 64.94 | 4.59E-08 |
| ncRNA export from nucleus | 64.94 | 1.21E-02 |
| SRP-dependent cotranslational protein targeting to membrane | 43.69 | 9.85E-33 |



**Figure S1. Related to Figure 1. MAGE-B2 regulates SG dynamics.**

(A) Tandem affinity purification coupled to mass spectrometry. HEK293 cells stably expressing vector control or TAP-MAGE-B2 were subjected to tandem affinity purification followed by LC-

MS/MS (n = 2). Gene ontology analysis of the identified proteins from mass spectrometry was performed using the PANTHER (protein annotation through evolutionary relationship) classification system according to biological process. The top enriched biological processes are ranked by fold enrichment over the expected value based on the reference list with raw *p* value determined by Fisher's exact test.
(B) MAGE-B2 knockdown does not cause spontaneous SG formation. U2OS cells were transfected with the indicated siRNAs for 72 hr and immunostained for G3BP1 and MAGE-B2. Representative images are shown.
(C and D) MAGE-B2 knockdown increases SG in HCT116 cells. HCT116 cells were transfected with the indicated siRNAs for 72 hr, treated with 62 μM sodium arsenite for 1 hr, and immunostained for G3BP1. Representative images shown in (C) and quantification (n = 3) of SG numbers per cell shown in (D). *P* values were determined by one-way ANOVA.
(E) WT or MAGE-B2 KO U2OS cells were treated with 62 μM sodium arsenite for 1 hr and immunostained for PABP-C1. Quantification (n = 3) of SG numbers per cell is shown. *P* values were determined by *t* test.
(F) WT, MAGE-B2 KO, or MAGE-B2-reconstituted KO U2OS cells were treated with 62 μM sodium arsenite for 1 hr and immunostained for PABP-C1. Representative images are shown. SG quantification is shown in Figure 1I.
(G) Overexpression of MAGE-B2 decreases SG. U2OS cells were transiently transfected with HA-MAGE-B2 for 48 hr, treated with 500 μM sodium arsenite for 1 hr, and immunostained for HA or G3BP1 (n = 3). Representative images are shown. Quantification of SG numbers per cell shown in Figure 1J.
(H and I) Live-cell imaging of U2OS cells stably expressing G3BP1-GFP. 72 hr after transfection with the indicated siRNAs, cells were treated with 500 μM sodium arsenite to induce stress granule assembly. Quantification of the percentage of SG-containing cells over time shown in (H). Quantification of the SG initiation time for each cell is shown in (I). Data from one representative experiment of n = 3 biological replicates with at least 40 cells analyzed for each group. *P* value was determined by unpaired *t* test.
(J and K) Live-cell imaging of U2OS cell stably expressing G3BP1-GFP. 72 hr after transfection with the indicated siRNAs, cells were treated with 1 μM thapsigargin to induce SG assembly. Quantification of the percentage of SG-containing cells over time is shown in (J). Quantification of the SG initiation time for each cell is shown in (K). Data from one representative experiment of n = 3 biological replicates with at least 20 cells analyzed for each group. *P* value was determined by unpaired *t* test.
(L) MAGE-B2 KO cells have similar growth rates to WT U2OS cells. WT, MAGE-B2 KO, or MAGE-B2-reconstituted KO U2OS cells were counted 72 hr after plating to determine relative growth rates. *P* value was determined by *t* test.
Data are mean ± SD. Asterisks indicate significant differences from the control (** = p ≤ 0.01, *** = p ≤ 0.001, **** = p ≤ 0.0001, n.s. = not significant).

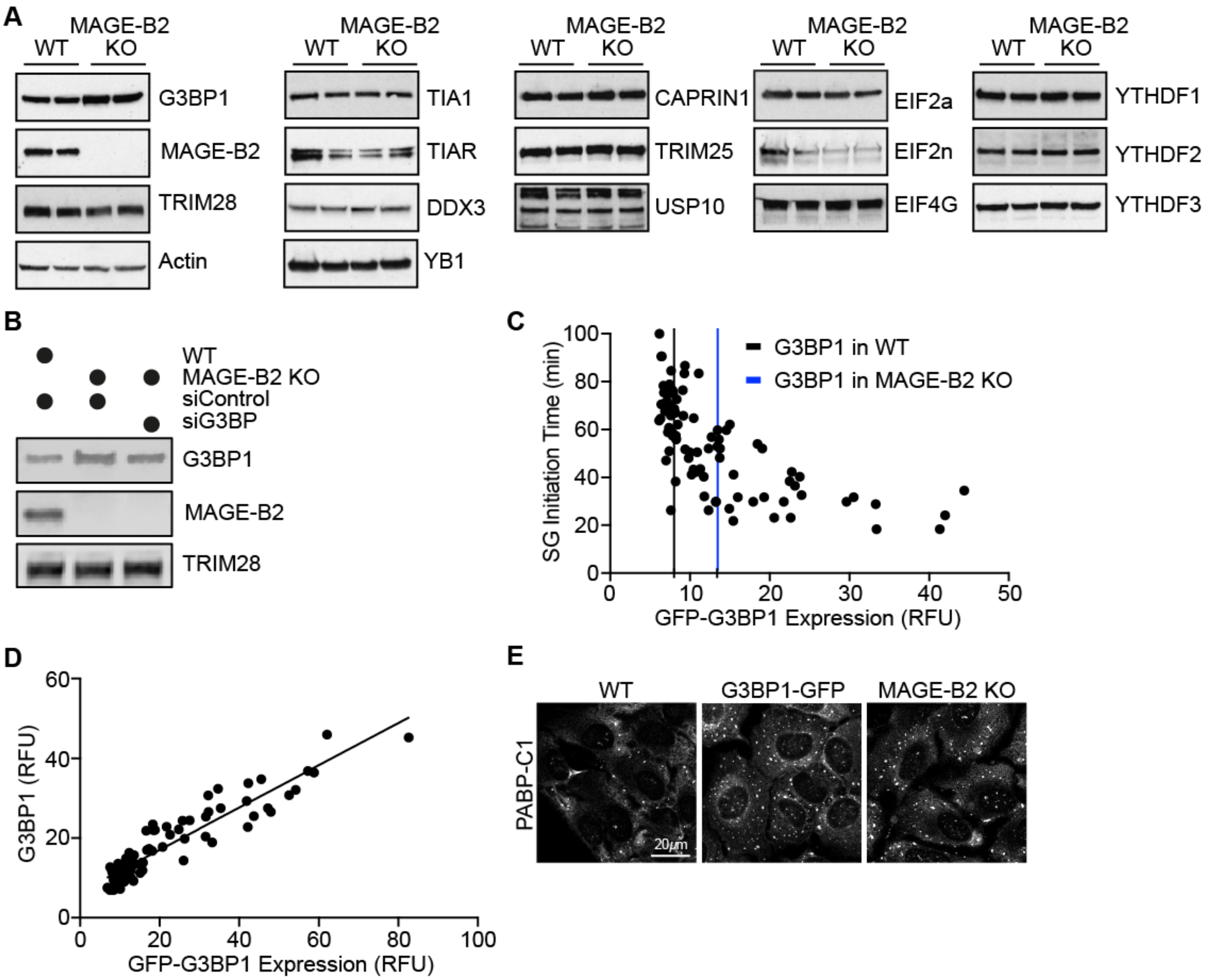


**Figure S2. Related to Figure 2. MAGE-B2-mediated regulation of G3BP protein levels results in altered SG dynamics.**

(A) MAGE-B2 regulates the protein levels of G3BP, but not that of other SG-associated proteins. WT or MAGE-B2 KO U2OS cell lysates were immunoblotted for the indicated proteins. Two biological replicate samples for each condition are shown.

(B) Rescue of G3BP protein levels in MAGE-B2 KO cells. WT or MAGE-B2 KO U2OS cells were transfected with the indicated siRNAs for 72 hr and immunoblotted for the indicated proteins.

(C) G3BP1 expression correlates with SG initiation time. G3BP KO U2OS cells were transiently transfected with varying levels of GFP-G3BP1. GFP-G3BP1 protein levels were determined at single-cell resolution by live-cell imaging before stress. Cells were then treated with 62 μM sodium arsenite to induce stress granule assembly (n = 3). The time at which SG assembly initiated was quantified and correlated to GFP-G3BP1 protein levels. Endogenous G3BP1 protein levels are indicated for WT or MAGE-B2 KO U2OS cells.

(D) Correlation between GFP-G3BP1 fluorescent intensity from GFP and immunofluorescent intensity from G3BP1 antibody staining. G3BP KO cells were transfected with GFP-G3BP1. 48 hr post-transfection, GFP intensity was determined at single-cell resolution by live-cell imaging. Cells were then immunostained for G3BP1 and fluorescence from the G3BP1 immunostaining was correlated to the previously measured GFP intensity on a cell-by-cell basis (n = 3, total 76 cells analyzed).

(E) Overexpression of G3BP1 is sufficient to increase SG formation similarly to MAGE-B2 KO. U2OS cells stably expressing G3BP1 were treated with 62 μM sodium arsenite for 1 hr and immunostained for PABP-C1. Representative images are shown. Quantification (n = 3) of SG numbers per cell shown in Figure 2K.

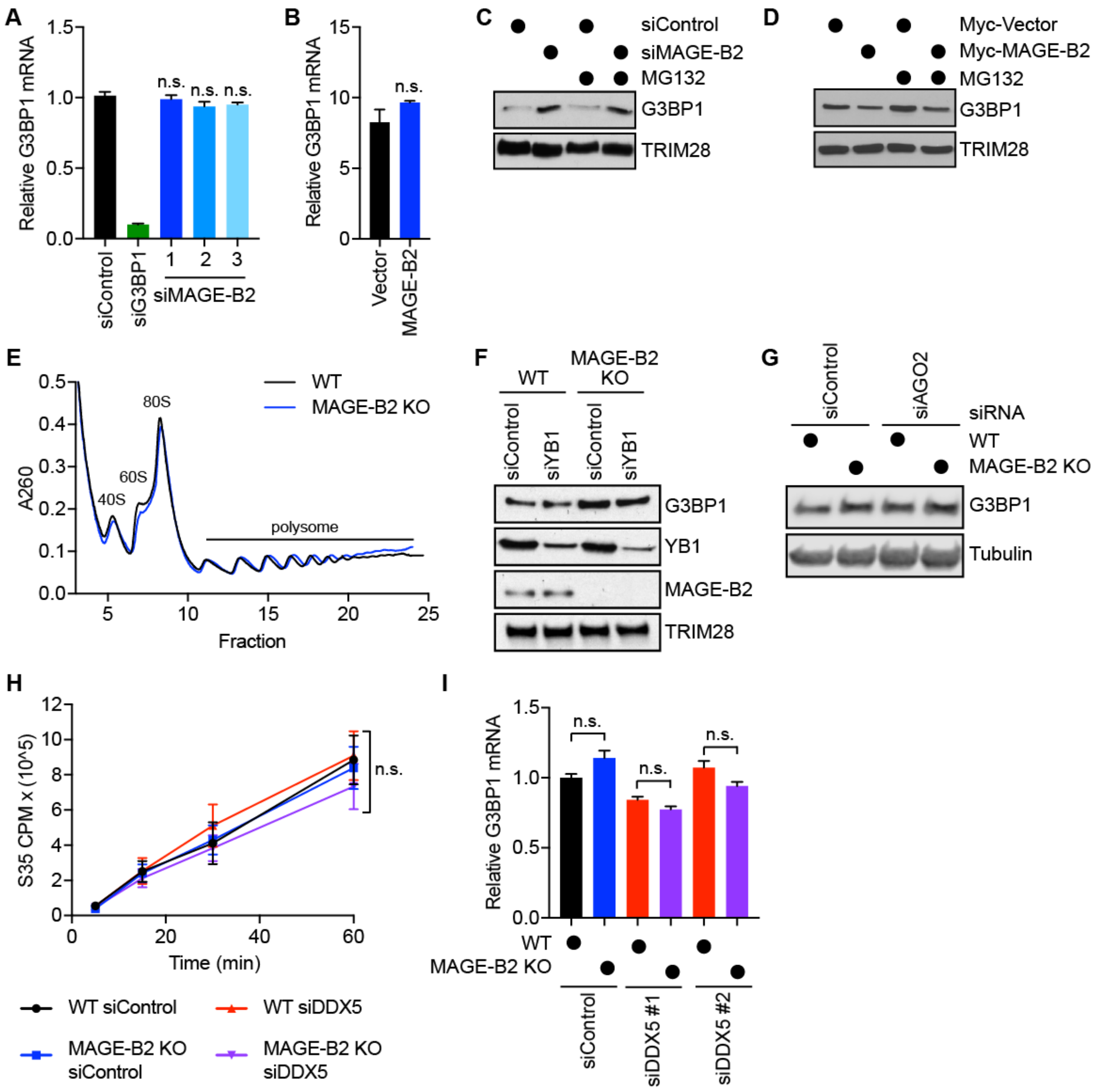


**Figure S3. Related to Figure 3. MAGE-B2 suppresses G3BP1 translation via inhibition of DDX5.**

(A) Knockdown of MAGE-B2 does not affect *G3BP1* mRNA levels. RT-qPCR analysis of G3BP1 mRNA levels normalized to 18s rRNA in U2OS cells transfected with the indicated siRNAs for 72 hr (n = 3). *P* values were determined by one-way ANOVA.

(B) Overexpression of MAGE-B2 does not affect *G3BP1* mRNA levels. RT-qPCR analysis of *G3BP1* mRNA levels normalized to *18S* rRNA in HeLa cells transiently transfected with either Myc-vector control or Myc-MAGE-B2 for 48 hr (n = 3). *P* value was determined by unpaired *t* test.

(C) Proteasomal inhibition does not affect G3BP1 protein levels. U2OS cells were transfected with the indicated siRNAs for 72 hr, treated with either DMSO or 10 μM MG132 for 6 hr, and immunoblotted for the indicated proteins.

(D) MAGE-B2 does not affect G3BP1 protein stability. HeLa cells were transfected with either Myc-vector control or Myc-MAGE-B2 for 48 hr, treated with either DMSO or 10 μM MG132 for 6 hr, and immunoblotted for the indicated proteins.
(E) MAGE-B2 KO does not affect global translation or polysome assembly. WT or MAGE-B2 KO U2OS cells were subjected to polysome fractionation by sucrose gradient. A260 nm was recorded during fraction collection. Data from one representative experiment of n = 4 biological replicates is shown.
(F) YB1 knockdown does not affect G3BP1 protein levels. WT or MAGE-B2 KO cells were transfected with control or YB1 siRNAs for 72 hr and immunoblotted for the indicated proteins.
(G) AGO2 knockdown does not affect G3BP1 protein levels. WT or MAGE-B2 KO cells were transfected with control or AGO2 siRNAs for 72 hr and immunoblotted for the indicated proteins.
(H) DDX5 knockdown does not affect global translation. WT or MAGE-B2 KO U2OS cells were transfected with the indicated siRNAs for 72 hr before global translation was measured by $^{35}$S-Met/Cys labeling for the indicated times and scintillation counting (n = 3). *P* values were determined by two-way ANOVA.
(I) DDX5 knockdown does not affect *G3BP1* mRNA levels. RT-qPCR analysis of *G3BP1* mRNA levels normalized to *18S* rRNA in WT or MAGE-B2 KO U2OS cells after transfection with the indicated siRNAs for 72 hr (n = 3). *P* values were determined by *t* test.
Data are mean ± SD. Asterisks indicate significant differences from the control (n.s. = not significant).

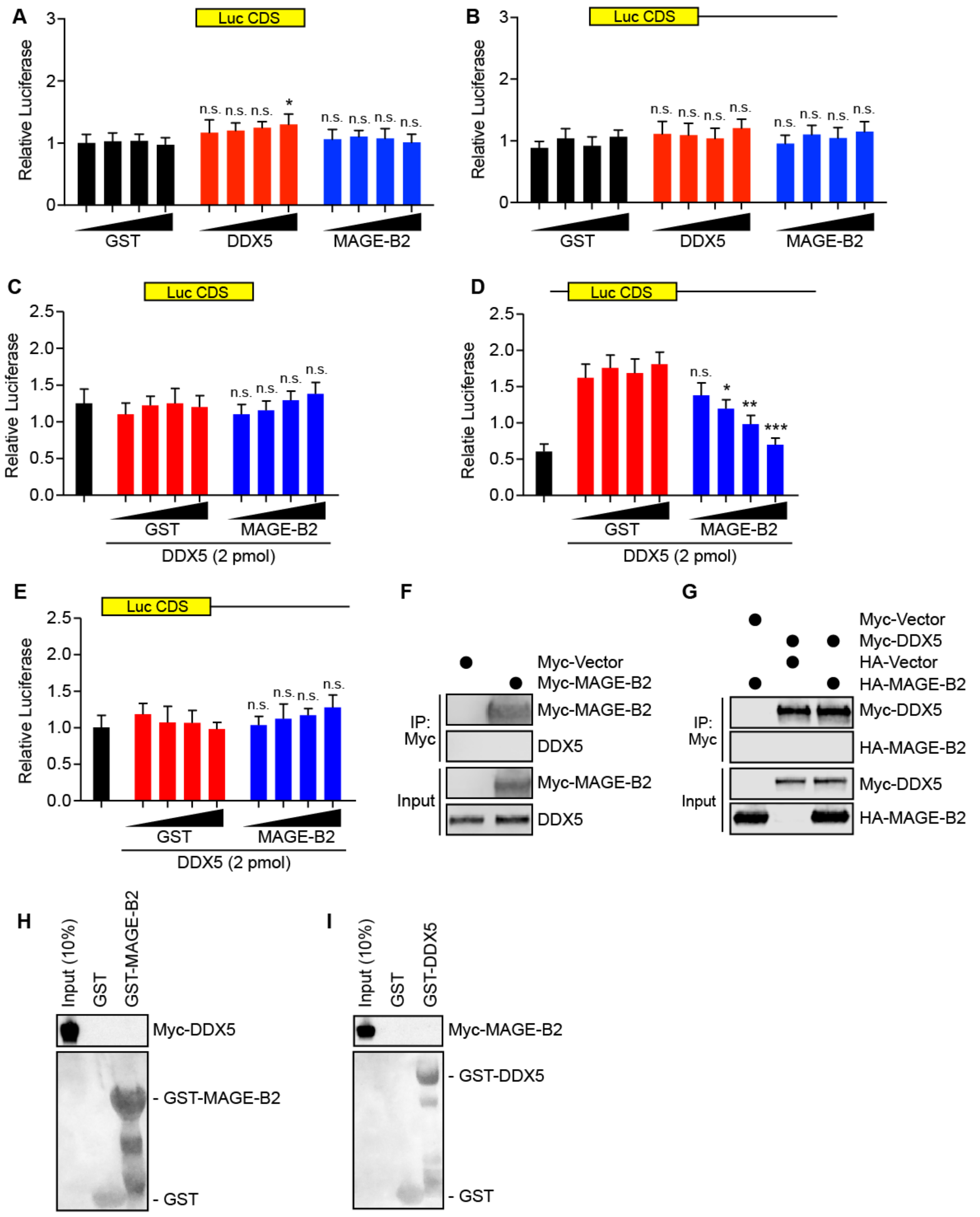


**Figure S4. Related to Figure 4. MAGE-B2 and DDX5 have opposing roles in the regulation of G3BP1 translation.**

(A) DDX5 and MAGE-B2 do not affect translation of luciferase without *G3BP1* UTRs. *In vitro* translation assays were performed using rabbit reticulocyte lysate with increasing amounts of recombinant GST (control), DDX5, or MAGE-B2 to measure translation of a control luciferase reporter (n = 3).

(B) DDX5 and MAGE-B2 do not regulate G3BP1 translation via the 3'UTR. *In vitro* translation assays were performed as in Figure S4A to measure translation of a luciferase reporter containing *G3BP1* 3'UTR (n = 3).

(C-E) *In vitro* translation assays reveal competition between DDX5 and MAGE-B2 for G3BP1 regulation via the 5' UTR. *In vitro* translation assays were performed as in Figure 4I to measure translation of a control luciferase reporter (C), a luciferase reporter containing G3BP1 5' and 3' UTRs (D), or only *G3BP1* 3' UTR (E) (n = 3).

(F and G) MAGE-B2 and DDX5 do not interact in cells. HeLa cells were transfected with the indicated constructs for 48 hr and immunoprecipitated with anti-Myc beads followed by immunoblotting for endogenous DDX5 (F) or overexpressed HA-MAGE-B2 (G).

(H and I) MAGE-B2 and DDX5 do not bind *in vitro.* Recombinant GST-MAGE-B2 (H) or GST-DDX5 (I) were incubated with *in vitro* translated Myc-DDX5 (H) or Myc-MAGE-B2 (I), pulled down by glutathione sepharose beads, eluted, separated by SDS-PAGE, and immunoblotted.

Data are mean ± SD. Asterisks indicate significant differences from the control (*p* values were determined by *t* test; * = $p \leq 0.05$, ** = $p \leq 0.01$, *** = $p \leq 0.001$, n.s. = not significant).

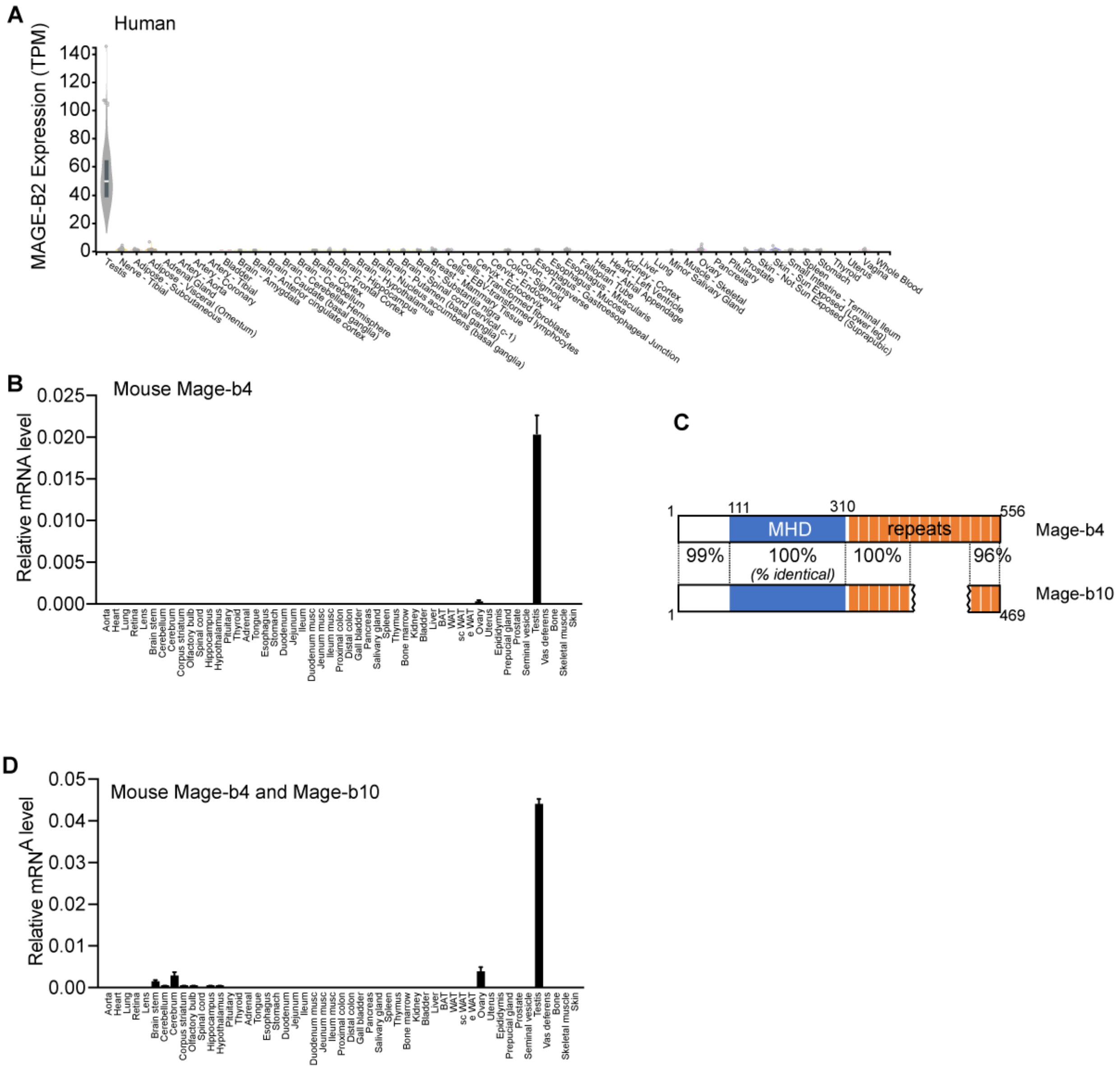


**Figure S5. Related to Figure 5. The mouse ortholog of human MAGE-B2 (Mage-b4) regulates stemness of testis spermatogonial stem cells.**

(A) Human *MAGE-B2* is expressed specifically in the testis. Gene expression data based on TPM values from the Genome-Tissue Expression (GTEx) Portal.

(B) Mouse *Mage-b4* is expressed specifically testis. RT-qPCR analysis (n = 3) of normalized mouse *Mage-b4* expression in the indicated tissues from C57BL/6 mice. Data are mean ± SD.

(C) Domain map and sequence identities of mouse Mage-b4 (top) and mouse Mage-b10 (bottom).

(D) Mouse *Mage-b10* is highly enriched in the testis. RT-qPCR analysis (n = 3) of normalized mouse *Mage-b4* and *Mage-b10* expression in the indicated tissues from C57BL/6 mice. Note RT-qPCR primer detects both *Mage-b4* and *Mage-b10* paralogs. Data are mean ± SD.

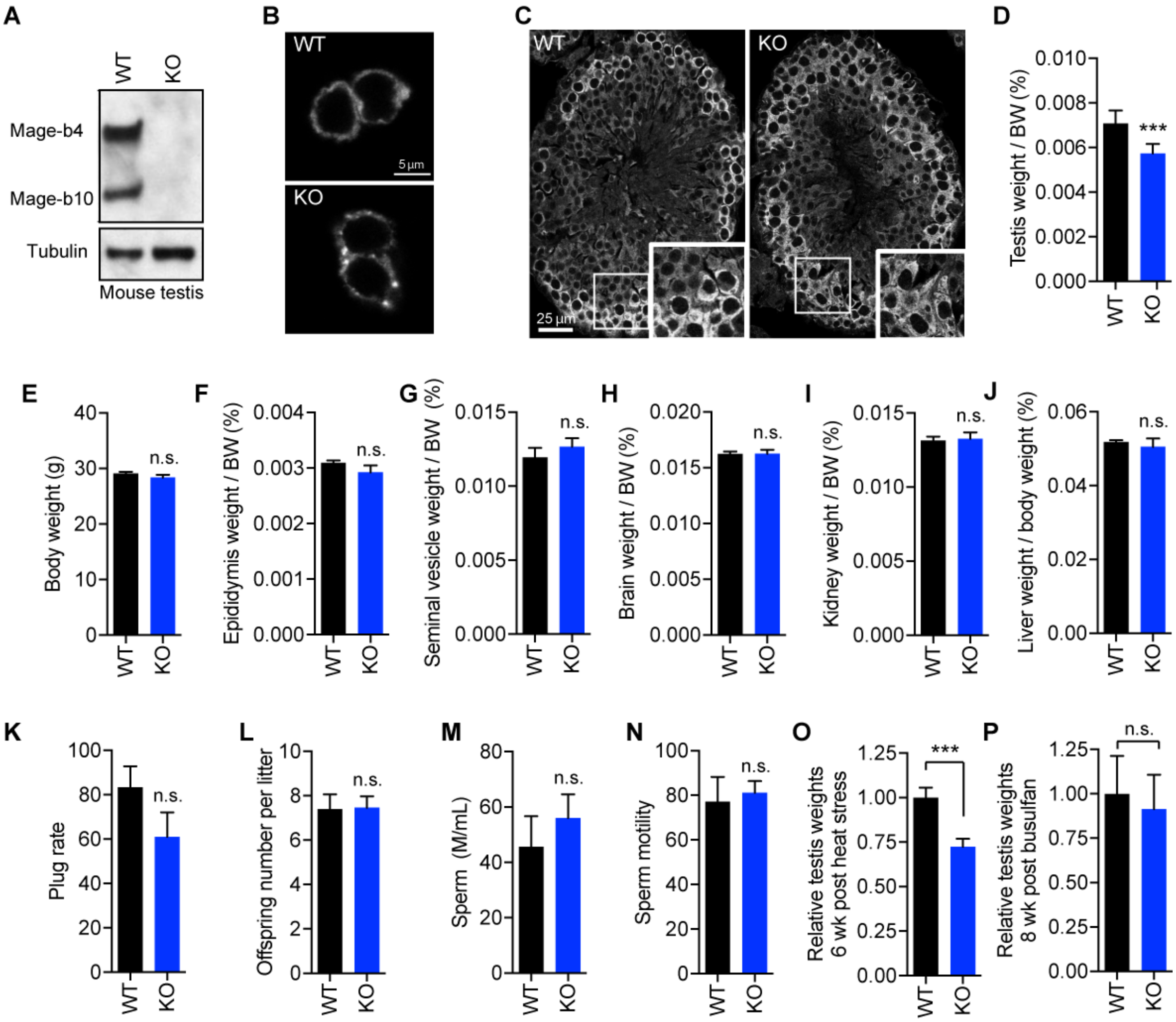


**Figure S6. Related to Figure 6. Knockout of the MAGE-B2 ortholog in mice results in hypersensitivity of the testis to heat stress.**

(A) Mage-b4 and its paralog Mage-b10 were knocked out by CRISPR/Cas9. Immunoblotting of testis lysates confirmed loss of Mage-b4/b10 protein.

(B) Primary cultures of undifferentiated spermatogonia from WT or Mage-b4 KO mice were treated with 250 μM sodium arsenite for 1 hr and immunostained. Representative images shown.

(C) Testis of WT or Mage-b4 KO mice were control stressed for 15 min at 33 °C. Mice were immediately sacrificed, testis isolated and sectioned before staining G3BP1 to detect stress granules. Representative images are shown.

(D-J) General characterization of Mage-b4 KO mice. Testis weights (D), body weights (E), epididymis weights (F), seminal vesicle weights (G), brain weights (H), kidney weights (I), and liver weights (J) are shown (n = 6 mice per genotype).

(K-N) Mage-b4 is not required for basal male fertility as shown by vaginal plug rates (K), number of pups born (L), sperm concentration (M), or sperm motility (N) compared to WT mice (n = 3-6 mice per genotype).

(O) Mage-b4 KO mice are hypersensitive to heat stress. WT or KO mice were heat shocked at 42 °C for 15 min and allowed to recover for 6 weeks before testis weights were measured (n = 6 per genotype). Testis weights were reported relative to WT mice and normalized to unstressed mice from each genotype.

(P) Mice were treated with busulfan as described in Figure 6J. Testis weights were measured 8 weeks post-treatment and normalized to untreated mice (n = 4-6 busulfan-treated mice per genotype, n = 8-10 control mice per genotype).

Data are mean ± SD. Asterisks indicate significant differences from the control (*p* values were determined by unpaired *t* test; *** = $p \leq 0.001$, n.s. = not significant).
